# A structurally unique effector shared between *Verticillium dahliae* and *Fusarium oxysporum* is involved in cotton defoliation and virulence on other hosts

**DOI:** 10.64898/2026.02.12.705609

**Authors:** Andrea Doddi, Gabriel Lorencini Fiorin, Jinling Li, Ilaria Zannini, Yukiyo Sato, Giulia Lancia, Giacomo Giuliari, Carmen Gómez-Lama Cabanás, Antonio Valverde-Corredor, Tingli Liu, Hui Tian, Grardy C.M. van den Berg, Jesús Mercado-Blanco, Baolong Zhang, Michael F. Seidl, Martijn Rep, Wessel Groot, Maria Carmela Bonaccorsi Di Patti, Francesca Troilo, Adele Di Matteo, Giorgio Giardina, Massimo Reverberi, Longfu Zhu, Luigi Faino, Bart P.H.J. Thomma

**Author notes:** Present address: Nanjing Engineering Research Center for Peanut Genetic Engineering Breeding and Industrialization, School of Food Science, Nanjing Xiaozhuang University, Nanjing 211171, China (Tingli Liu). Present address: Department of Soil and Plant Microbiology, Zaidín Experimental Station, CSIC, Profesor Albareda 1, 18008 Granada, Spain. These authors contributed equally (Andrea Doddi, Gabriel Lorencini Fiorin, Jinling Li, Ilaria Zannini, Yukiyo Sato). These authors contributed equally (Luigi Faino, Bart P.H.J. Thomma). To whom correspondence should be addressed., &.

## Abstract

Defoliating (D) strains of the vascular wilt fungus *Verticillium dahliae* cause severe yield losses in cotton and olive worldwide, yet the genetic basis underlying this pathotype has remained unknown. Here, we combined comparative genomics, functional genetics, structural analysis, and phylogenomics to uncover the molecular determinant of defoliation. We identified a small, D pathotype–specific genomic region encoding two duplicated secreted effector genes. Simultaneous deletion of both gene copies abolished pathogenicity and defoliation in cotton, olive, *Nicotiana benthamiana*, and *Arabidopsis thaliana*, whereas single deletions reduced virulence and genetic complementation restored disease symptoms. Conversely, expression of the D effector in non-defoliating strains was sufficient to induce defoliation. Moreover, or exogenous application of the purified protein, induced wilting and defoliation as well. Structural analyses revealed that D homologs share a conserved but previously uncharacterized protein fold and are distributed across *Verticillium* and *Fusarium* species, exhibiting functional diversification and host-specific activity. Phylogenomic and genomic context analyses indicate repeated horizontal transfer events mediated by giant transposable elements known as *Starships*. Together, our findings identify the D effector as a central pathogenicity factor that drives defoliation and virulence, and demonstrate how *Starship*-mediated horizontal gene transfer shapes the emergence and dissemination of an agriculturally devastating fungal trait.

## INTRODUCTION

*Verticillium dahliae* is a soil-borne fungal pathogen that causes vascular wilt disease by colonizing the water-conducting xylem vessels of hundreds of dicotyledonous host plants, including cotton (*Gossypium hirsutum*), olive (*Olea europaea*), and tomato (*Solanum lycopersicum*)^1,2^. Infection typically starts once microsclerotia that reside in the soil germinate, triggered by root exudates, and the fungus penetrates the root. Subsequently, the fungus enters xylem vessels where further proliferation takes place, affecting water transport, and causing characteristic symptoms that include wilting, stunting, chlorosis and early senescence^2,3^. Due to its broad host range, long-term prevalence of its resilient survival structures in the soil, limited curative treatments and scarcity of disease resistance in crop germplasm, Verticillium wilt control is notoriously difficult^2,4^

Although *V. dahliae* causes wilt disease in a broad range of host plants, virulence capacities and the severity of symptoms that are induced on host plants can vary considerably between individual strains. For instance, particular strains of *V. dahliae* can cause severe symptoms that include defoliation on cotton, olive, okra (*Hibiscus esculentus*) and pistachio (*Pistacia vera*), while other strains cause milder wilting symptoms and do not cause defoliation^5–8^. Consequently, on particular host plants, *V. dahliae* strains that show differential virulence capacities and disease symptoms are assigned to specific “pathotypes”^5,6^. Thus, *V. dahliae* strains that are highly virulent and cause rapid and severe defoliation on cotton, olive, okra and pistachio are assigned the defoliating (D) pathotype, whereas strains that are moderately virulent and only induce mild wilting symptoms without defoliation are assigned to the non-defoliating (ND) pathotype^5^. The currently increasing prevalence of the highly virulent D pathotype strains poses a significant threat to cotton and olive plantations worldwide^9,10^. It is believed that D pathotype strains originated once in North America and subsequently spread to other continents through the dispersal of contaminated plant commodities^11,12^. Despite many efforts to develop molecular markers to robustly and accurately differentiate D and ND pathotype strains^13–15^, the molecular basis for the difference in aggressiveness between D and ND pathotype strains remains controversial so far.

To establish disease on their host plants, adapted pathogens secrete so-called effector molecules, many of which will modulate host physiology, including host immune responses, to promote successful infection^16,17^. However, for several pathogens, including *V. dahliae,* it was recently shown that particular effectors target host-associated microbiota to promote disease development^18–22^. Thus, it is well established that comprehensive identification of effector repertoires and determination of their modes of action is key to uncover virulence mechanisms and infection strategies of plant pathogens, which is ultimately required to design effective disease management strategies ^23^. Here, we aimed to elucidate the molecular basis for the differential aggressiveness between *V. dahliae* strains that belong to the D and ND pathotype, respectively.

## RESULTS

### Comparative genomics identifies D pathotype-specific effector genes

A diverse set of *V. dahliae* strains had previously been sequenced, including well-characterized D pathotype strains (V991, T9, V117, and TM6) and ND pathotype strains (BP2, V4, cd3, and HN)^5,10,24–26^. To expand the set of strains for comparative genomics aimed at identifying determinants of defoliation, we generated a phylogenetic tree of in-house sequenced strains (Table S1; Figure S1). This analysis identified six additional strains clustering with known D pathotype strains, while others grouped with ND strains. Four strains (V574, V700, V679, and Vd39) clustered with D strains but were phylogenetically distinct, prompting disease assays to determine their pathotype. We also included presumed D strains (V76, V117, CQ2, ST100) and presumed ND strains (JR2, VdLs17). Disease assays confirmed V76, V117, CQ2, and ST100 as D pathotype strains, whereas JR2, VdLs17, V574, V700, and Vd39 were classified as ND (Figure 1).

**Figure 1.**
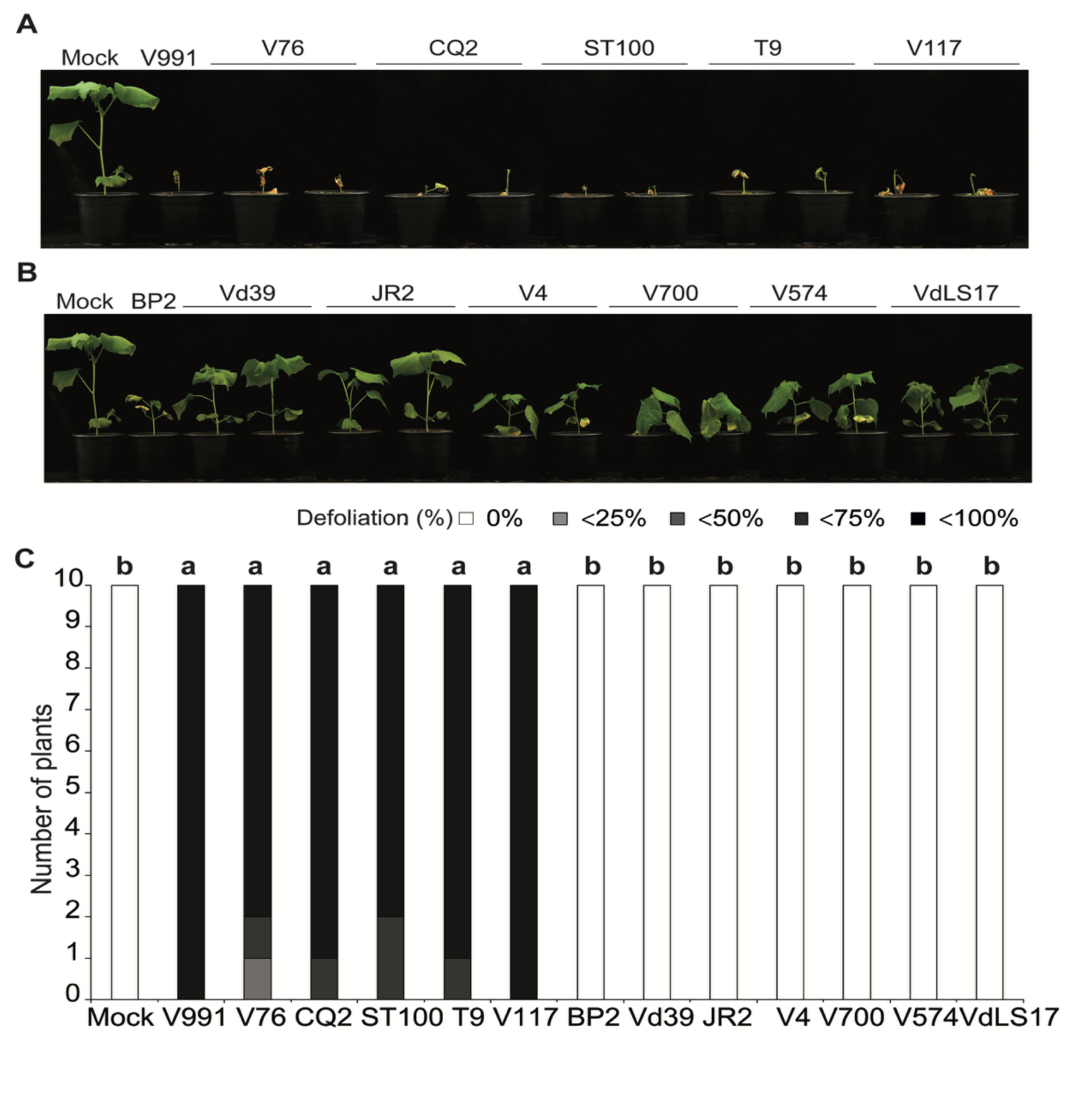
Phenotype of cotton plants inoculated with various sequenced *Verticillium dahliae* strains. (A) Typical phenotype of cotton plants (cv. Xinluzao63) upon mock-inoculation or inoculation with *V. dahliae* strains V991, V76, CQ2, ST100, T9 and V117 at 28 days post inoculation (dpi). Previously characterized D pathotype strain V991 used as inoculation control. (B) Typical phenotype of cotton plants upon mock-inoculation or inoculation with *V. dahliae* strains BP2, Vd39, JR2, V4, V574, V700, VdLS17 at 28 dpi. Previously characterized ND pathotype strain BP2 used as inoculation control. (C) Defoliation was classified as 0 (0% leaf drop off), 1 (<25% leaf drop off), 2 (<50% leaf drop off), 3 (<75% leaf drop off) and 4 (<100% leaf drop off) at 28 dpi. Inoculation experiments were performed with ten plants for each fungal strain and repeated twice independently with similar results.

The genome of the D pathotype strain CQ2 was sequenced using PacBio (GCA_004798895.1) and used as the reference for comparative genomics with eight D pathotype strains (V991, T9, V117, TM6, V76, 463, CQ2, and ST100) and nine ND pathotype strains (BP2, V4, HN, cd3, JR2, VdLs17, V574, V700, and Vd39)^27,28^. This analysis identified ∼200 kb of D-specific sequence harboring ∼30 predicted protein-coding genes. To further refine the candidate region, genomes of 72 additional *V. dahliae* strains were analyzed (Figure S2). One strain, BGI_32, clustered outside both D and ND groups and was subsequently classified as ND based on disease assays (Figure S3), allowing the D-specific interval to be reduced to ∼24 kb containing seven genes. InterProScan analysis^29^ showed that only two genes (CQ2_Uni028g00060 and CQ2_Uni008g00030) encode predicted secreted proteins and are thus candidate effectors. Transcriptome analysis of cotton plants inoculated with D pathotype strain V991 revealed high expression of both genes (Figure S4). Notably, the two genes are 100% identical despite residing on separate contigs (Unitig_28 and Unitig_8), and alignment of these contigs showed identical flanking regions, indicating segmental duplication (Figure 2). These *in planta* expressed genes were tentatively named *D* genes due to their potential role in cotton defoliation.

**Figure 2.**
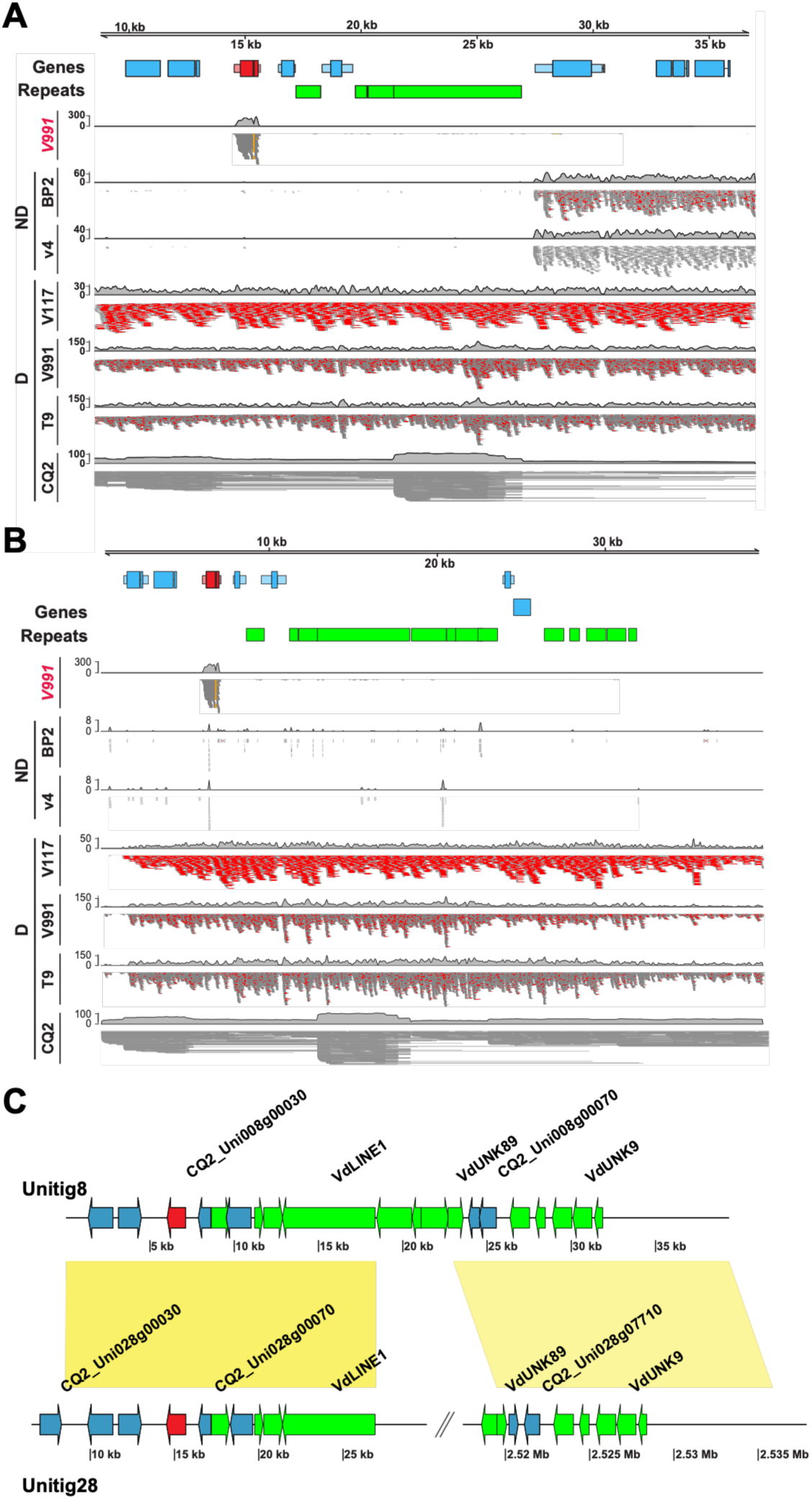
Two identical copies of effector gene generated by segmental duplication. (A-B) Schematic representation of genomic region (Unitig_28 and Unitig_8) where two candidate effector genes located. Gene models shown in blue, the candidate effector genes displayed in red, while repetitive elements displayed in green. RNA reads from cotton plants infected by a D pathotype strain (V991 in red) that only mapped to candidate gene are showed in dark grey. Reads from DNA-seq of ND pathotype strains (V4 and BP2) and D pathotype strains (V117, V991 and T9) mapped onto CQ2 genome. The reads are showed as coverage plot and the depth of DNA reads coverage indicated in red. (C) Syntenic assignment of Unitig_28 and Unitig_8, the extensive synteny (yellow blocks indicate 100% sequence identity) points towards a segmental duplication. Gene models shown in blue, the candidate effector gene displayed in red, while repetitive elements displayed in green.

### The D effector is responsible for cotton defoliation

To further investigate the role of the D effector in cotton pathogenicity, targeted deletion of the two *D* gene copies was performed in the D pathotype strain CQ2 by sequential homologous recombination (Figure S5A). Single-copy and double-copy deletion mutants (Δ*D* and ΔΔ*D*) were generated and verified by PCR (Figure S5B). Cotton seedlings were inoculated with wild-type CQ2, single deletion mutants (Δ*D*#1, Δ*D*#2), or double deletion mutants (ΔΔ*D*#1, ΔΔ*D*#2), and disease progression was monitored. Wild-type CQ2 induced severe defoliation at 28 days post inoculation (dpi) (Figure 3A–B), whereas Δ*D* mutants showed significantly reduced aggressiveness and defoliation. In contrast, ΔΔ*D* mutants failed to cause disease, with no defoliation symptoms and no detectable fungal biomass by quantitative PCR, demonstrating that *D* is required for pathogenicity (Figure 3A–C). Complementation of single and double deletion mutants with a genomic fragment containing the D coding sequence (*pD::D*) restored aggressiveness and defoliation (Figure 3; Figure S6). Introduction of *pD::D* into the ND pathotype strain JR2 increased aggressiveness on cotton, causing severe stunting and defoliation (Figure S7, S8), confirming the role of D as a pathogenicity factor of D pathotype strains.

**Figure 3.**
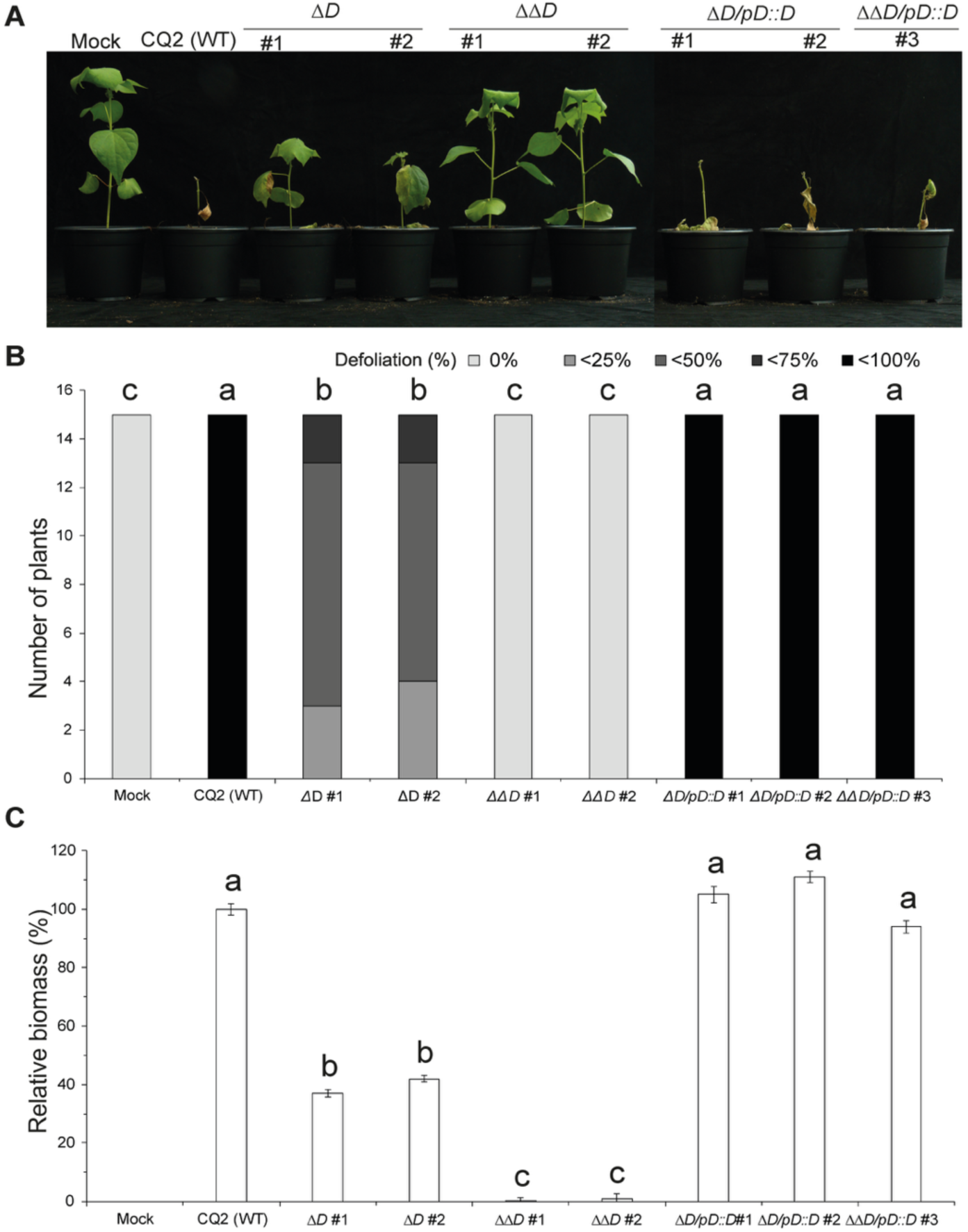
D effector is responsible for cotton defoliation. (A) Typical phenotype of cotton (cv. Xinluzao63) upon mock-inoculation or inoculation with wild type strain CQ2 (WT), two Δ*D* mutants (Δ*D* #1 and Δ*D* #2), two ΔΔ*D* mutants (ΔΔ*D* #1 and ΔΔ*D* #2), two Δ*D* complementary strains (Δ*D*/p*D*::*D* #1 and Δ*D*/p*D*::*D* #2) and one ΔΔ*D* complementary strain (ΔΔ*D*/p*D*::*D* #3) at 28 days post inoculation (dpi). (B) Defoliation were classified as 0 (0% leaf drop off), 1 (<25% leaf drop off), 2 (<50% leaf drop off), 3 (<75% leaf drop off) and 4 (<100% leaf drop off). (C) Fungal biomass as determined with real-time PCR at 28 dpi. Bars represent *V. dahliae* ITS levels relative to cotton ubiquitin (*Gh*UB) levels (for equilibration) with standard deviation in a sample of five pooled plants. The fungal biomass in cotton plants upon inoculation with the wild type CQ2 is set to 100%. Different letter labels indicate statistically significant differences (Student’s t-test; P < 0.05). Experiments were repeated twice independently with similar results.

To determine whether the D effector protein alone can induce Verticillium wilt symptoms, cotton seedlings were treated with heterologously produced D protein (9 μM) and monitored over time, with water and the chitin-binding effector Vd2LysM (9 μM) as negative controls^30^. D protein treatment induced wilting, mild chlorosis, and initial cotyledon detachment after 10 days, progressing to severe chlorosis and detachment of more than half of the leaves (55 ± 6.45%) after 16 days (Figure S9). No symptoms were observed in control treatments, indicating that the D effector protein itself, rather than extensive fungal colonization, is sufficient to cause defoliation symptoms.

### The D effector mediates olive defoliation and pathogenicity on *Nicotiana benthamiana* and *Arabidopsis thaliana*

Since the D effector was shown to mediate cotton defoliation, we examined whether olive-defoliating *V. dahliae* strains carry the *D* gene. The *D* gene was detected in previously characterized D pathotype strains V150I, V641I, V356I, and V403II^31,32^, but not in the ND pathotype olive strain 812I^31^ (Figure S10). To test whether D mediates olive defoliation, both *D* gene copies were deleted in strain V150I (Figure S5C), and olive plants were inoculated with three double-deletion mutants (ΔΔ*D*#5, ΔΔ*D*#7, ΔΔ*D*#8) or wild-type V150I. While wild-type V150I caused severe disease, including defoliation and plant death, ΔΔ*D* mutants showed greatly reduced pathogenicity (Table 1, Figure S11), demonstrating that the D effector is responsible for olive defoliation.

**Table 1.**
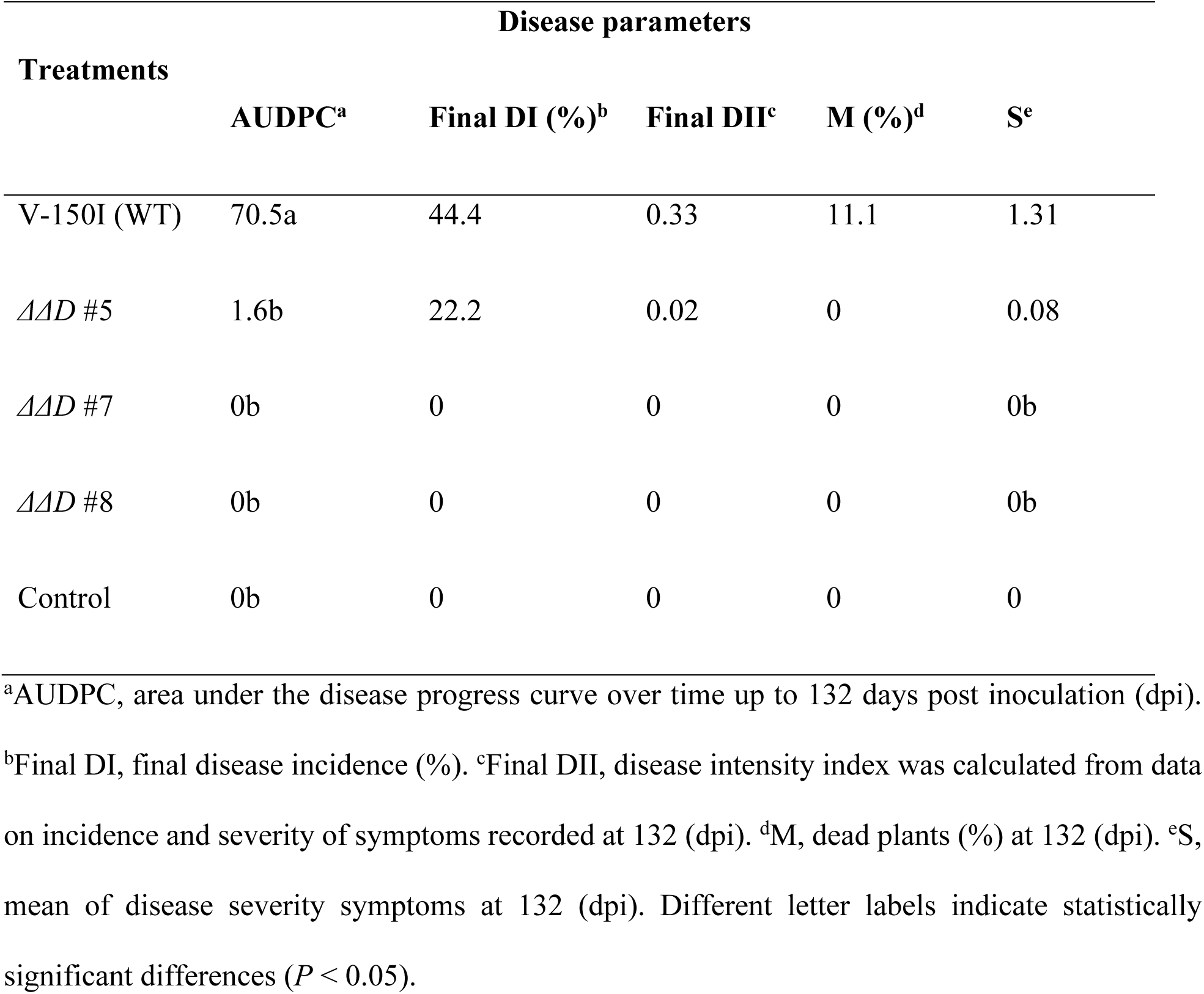
Bioassay of *D* gene deletion strains on olive (cv. Picual) plants.

To determine whether the role of the D effector extends beyond cotton and olive, Δ*D* and ΔΔ*D* mutants were inoculated on *Nicotiana benthamiana* and *Arabidopsis thaliana*. On *N. benthamiana*, Δ*D* mutants showed reduced aggressiveness, with increased canopy area and lower fungal biomass, while ΔΔ*D* mutants were non-pathogenic, showing no disease symptoms or detectable fungal biomass (Figure 4A–C). Similarly, on *A. thaliana*, Δ*D* mutants exhibited reduced aggressiveness, as indicated by larger rosette leaf area and lower fungal biomass, whereas ΔΔ*D* mutants were non-pathogenic (Figure 4D–F). These results indicate that the D effector is required not only for defoliation on highly susceptible hosts but also for general pathogenicity on *N. benthamiana* and *A. thaliana*.

**Figure 4.**
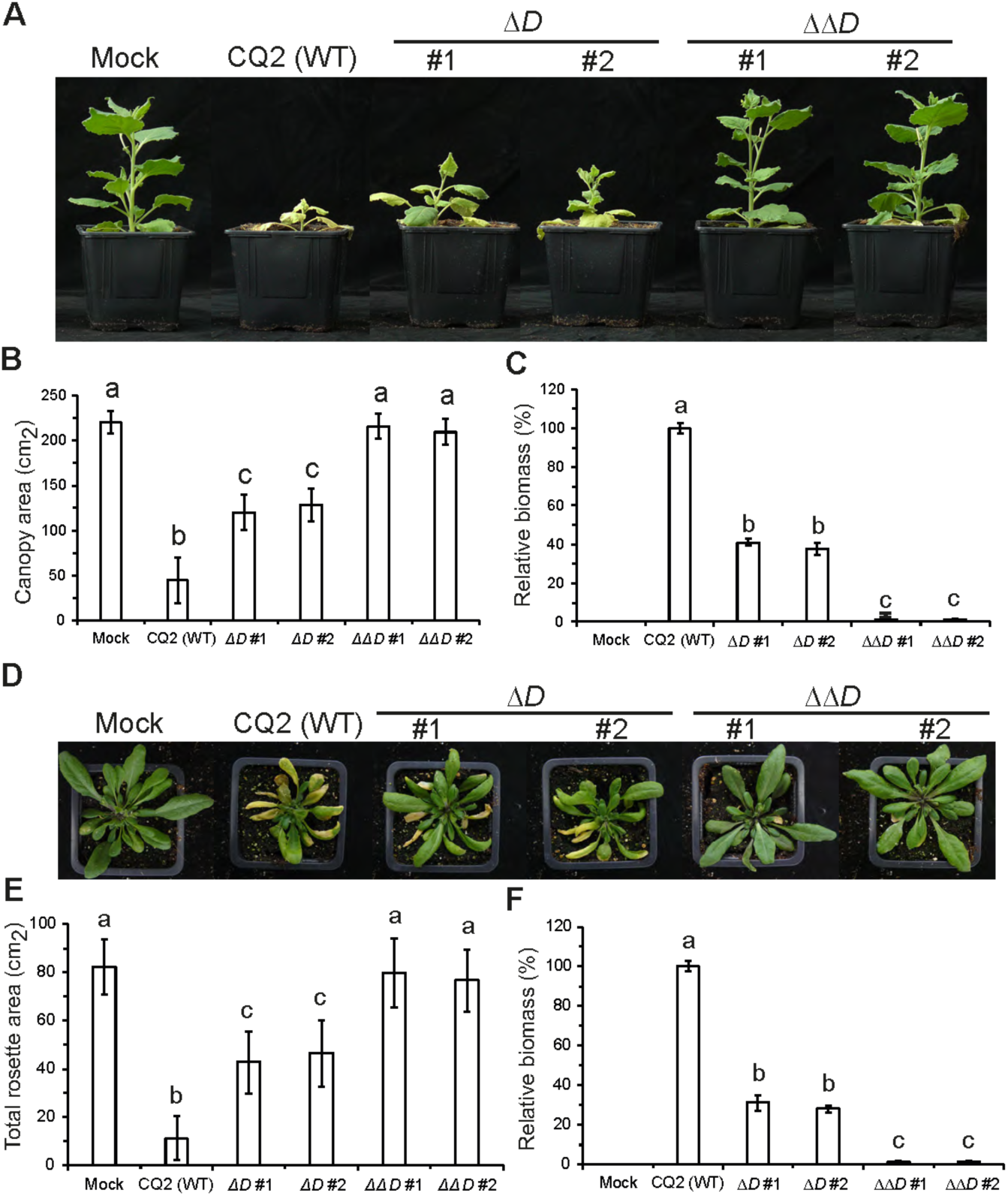
D effector is a pathogenicity factor on *Nicotiana benthamiana* and *Arabidopsis thaliana*. (A) Typical phenotype of *N. benthamiana* plants that were mock-inoculated or inoculated with wild type strain CQ2 (WT), two Δ*D* mutants (#1 and #2), two ΔΔ*D* mutants (#1 and #2) at 21 days post inoculation (dpi). (B) Quantification of the canopy area of *N. benthamiana* at 21 dpi. Bars represent the average canopy area of five plants with standard deviation. (C) Fungal biomass as determined with real-time PCR at 21 dpi. Bars represent *V. dahliae* ITS levels relative to *Nb*RuBisCo (for equilibration) with standard deviation in a sample of five pooled plants. The fungal biomass in *N. benthamiana* plants upon inoculation with the wild type CQ2 is set to 100%. Different letter labels indicate statistically significant differences (Student’s t-test; P < 0.05). (D) Typical phenotype of *A. thaliana* (Col-0) plants that were mock-inoculated or inoculated with indicated fungal strains in panel A at 21 (dpi). (E) Quantification of the rosette area of five *A. thaliana* plants at 21 dpi. Bars represent the average rosette area of five plants with standard deviation. (F) Fungal biomass as determined with real-time PCR at 21 dpi. Bars represent *V. dahliae* ITS levels relative to *At*RuBisCo (for equilibration) with standard deviation in a sample of five pooled plants. The fungal biomass in *A. thaliana* plants upon inoculation with the wild type CQ2 is set to 100%. Different letter labels indicate statistically significant differences (Student’s t-test; P < 0.05). *N. benthamiana* and *A. thaliana* inoculation experiments were performed with five plants for each fungal strain and repeated twice independently with similar results.

### D effector homologs occur in the *V. dahliae* population

To assess the distribution and variability of the D effector in *V. dahliae*, 63 genome sequences were queried for the presence of the *D* gene. Interestingly, while all D pathotype strains carry D homologs that are 100% identical, diverged homologs occur throughout the *V. dahliae* population encoding D homologs that display 91% to 82% amino acid similarity, corresponding to seven *D*-*like* alleles (*D-like1* to *D-like7*) (Figure 5A). *D-like1* was present in 22 strains, including reference genomes JR2 and VdLs17 ^33^; *D-like2* in 23 strains; *D-like3* in five strains; *D-like4* in two strains; *D-like5* in eleven strains; and *D-like6* and *D-like7* each in a single strain, I1V. *D-like1*, *D-like2*, and *D-like3* encode ∼237 amino acid proteins ∼80% identical to D and differ from each other by only one residue (Figure 5B). In contrast, *D-like4*, *D-like5*, and *D-like7* encode truncated 82-amino-acid proteins due to premature stop codons, with *D-like4* additionally disrupted by a *Ty3*/*Gypsy*-like retrotransposon in V574 and V700 (Figure S12). *D-like6* co-occurs with *D* on the same contig in strain I1V.

**Figure 5.**
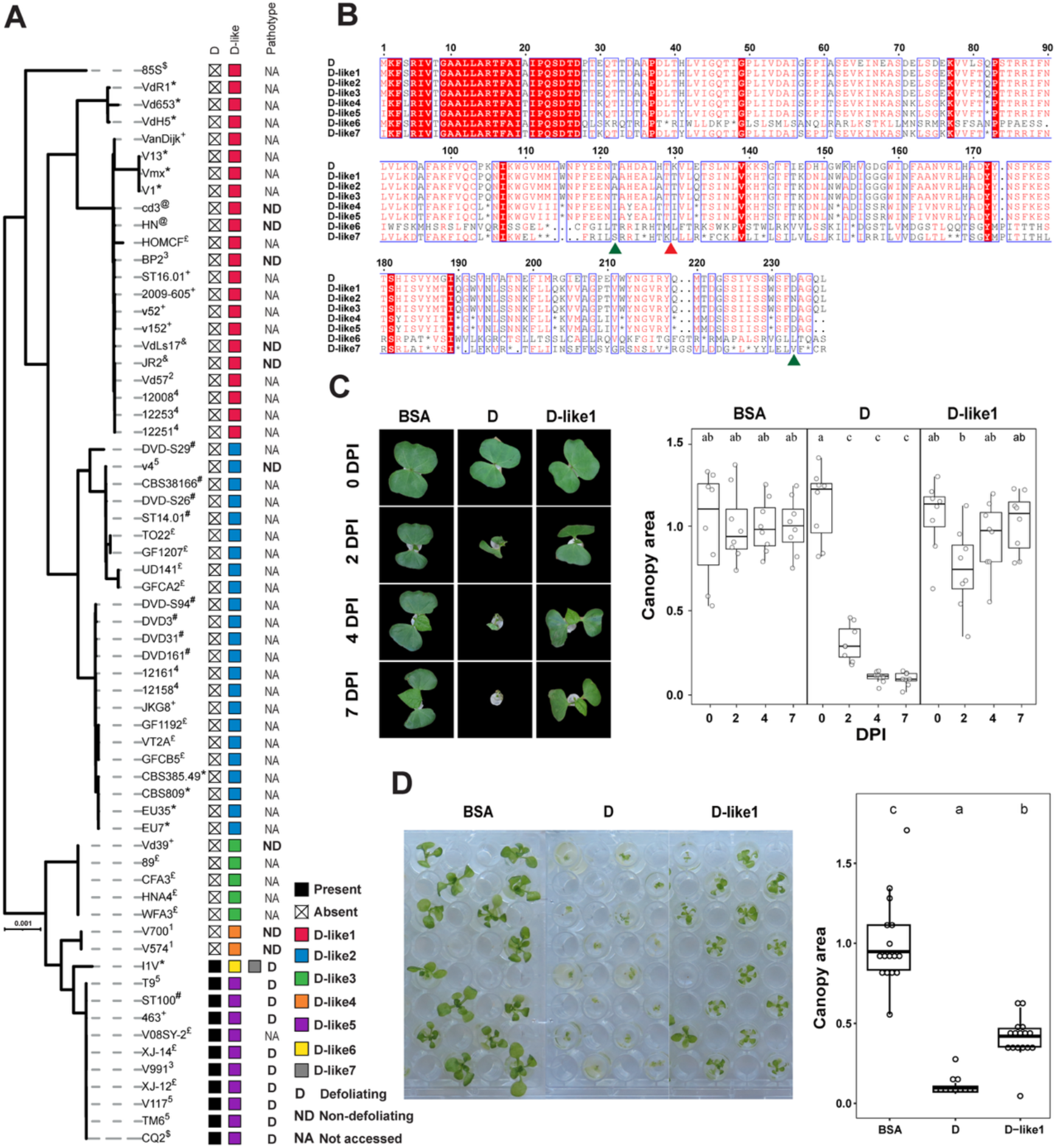
A multiallelic *D-like* gene occurs in the *V. dahliae* population. (A) Distribution of the *D* gene and *D-like* alleles in the *V. dahliae* population. Phylogenetic relationships among sequenced *V. dahliae* strains were determined based on whole genome alignments and branch lengths in the tree indicate sequence divergence. £= Chavarro-Carrero et al., 2021; * = Chavarro-Carrero et al., unpublished; @= Xu et al., 2012; $= Depotter et al., 2019; += Gibriel et al., 2019; &= Faino et al., 2015; #= de Jonge et al., 2012; 1= Milgroom et al., 2016; 2= Li, 2019; 3= Zhang et al., 2012; 4= Fan et al., 2018; 5= Keykhasaber, 2017 (B) Alignment of the predicted proteins encoded by the *D* gene and by each of the *D-like* alleles. The red arrowhead indicates the position at which the *D-like* 4 allele is disrupted by a retrotransposon insertion. The green arrowheads indicate the positions at which differences among the highly identical proteins encoded by the alleles *D-like* 1, *D-like* 2 and *D-like* 3 are observed. Asterisks indicate stop codons. The numbers on top of the alignment indicate amino acid positions in the sequences. (C) Typical appearance of cotton seedlings (cv. Xinluzao63) upon application of *D* and *D-like*1 effector proteins from *Verticillium dahliae* (5 µM), at 0, 2, 4 and 7 days post treatment. Bovine serum albumin (BSA; 5 µM) was used as control. Quantification of the canopy area of treatments shown is normalised by the average of BSA treated plants at 0, 2, 4 and 7 days after start of the treatment. Each bar represents eight plants. Different letter labels indicate statistically significant differences (one-way ANOVA and Tukey test). Experiments were repeated twice with similar results. (D) Typical appearance of *Arabidopsis thaliana* seedlings (Col-0) upon application of *D* and *D-like1* effector proteins from *Verticillium dahliae* (5 µM), at two weeks after seed germination. Bovine serum albumin (BSA; 5 µM) was used as a control. Strong chlorosis and growth arrest are visible. Quantification of the canopy area of treatments shown is normalised by the average of BSA-treated plants at two weeks after the start of the treatment. Each bar represents sixteen plants. Different letter labels indicate statistically significant differences (one-way ANOVA and Tukey test). Experiments were repeated twice with similar results.

To test functional activity, D-like1 protein was heterologously produced in *E. coli* and cotton seedlings were exposed to 5 μM D-like1, with D protein as a positive control. While D protein induced wilting, chlorosis, and >50% leaf detachment by 16 days (Figure 5C), D-like1 caused no symptoms, similar to BSA, indicating it cannot trigger cotton defoliation. *Arabidopsis thaliana* seedlings grown on 5 μM D protein exhibited chlorosis and arrested growth at the two-leaf stage, whereas D-like1 also caused significant growth inhibition relative to BSA-treated controls (Figure 5D). These results indicate that *D-like1* exerts differential effects depending on the plant species.

### *Starship* giant transposon-mediated *D* gene transfer between *Fusarium* and *Verticillium*

To determine whether D protein homologs occur beyond *V. dahliae*, ∼22,000 fungal genomes from NCBI were queried, identifying 19 additional homologs (Table S3). Within *Verticillium*, *D* homologs were found in *V. alfalfae*, and *V. longisporum*, but not in other species. *V. alfalfae* strains Ms102 and PD683 harbor a novel *D-like8*. *V. longisporum* strains showed complex patterns: vl1 has five copies (*D-like8*, *D-like10*, *D-like11*, and two *D-like9*), PD589 and VLB2 have three copies with two *D-like8* and differing third copies (*D-like12* in PD589, *D-like9* in VLB2), Vl43 contains *D-like8* and *D-like9*, and Vl32 carries *D-like1*.

Notably, outside *Verticillium*, *D* homologs were detected only in *Fusarium* species; two in *F. secorum* and 17 in ten *formae speciales* of *F. oxysporum*, with 80–90% identity to *V. dahliae* D (Table S3). Most strains carried a single homolog, but some had multiple copies. *F. oxysporum* f. sp. *vasinfectum* (*Fov*) and f. sp. *melongenae* (*Fomel*) had two identical copies, one *Fov* strain had three identical copies plus a fourth variant, and f. sp. *melonis* (*Fom*) had the highest diversity with four *D-like* versions (*D-like13*, *D-like14*, *D-like15*, *D-like27*). *D-like13*, present in 30 of 48 *Fom* strains, encodes a 27-aa peptide disrupted by an inserted gene. *Fom* strains R12/13 and NRRL26174 carry both *D-like15* and *D-like27* (Table S3).

The restricted occurrence of *D* homologs to *Verticillium* and *Fusarium*, which belong to the distant fungal orders Glomerellales and Hypocreales, suggests horizontal gene transfer (HGT). To test this, alien index^34^ (AI) scores were calculated for 12,882 Pezizomycotina genomes using non-*Verticillium* Glomerellales as ingroup and all other genera as outgroup. The *D* gene exhibited an AI score of 460.5, strongly supporting HGT across orders.

To investigate the divergence of *D* homologs, a phylogeny based on nucleotide sequences was inferred. *D* homologs from *Verticillium* formed a distinct clade from those in *Fusarium*, suggesting a single HGT event between the two genera (Figure S13A). To estimate its timing, synonymous substitution rates (*K_s_*) of the *D* gene were compared with the vertically inherited *GAPDH* gene. The *K_s_* between the *V. dahliae D* gene and its closest *Fusarium* ortholog was lower than *GAPDH K_s_* between *V. dahliae* and *Fusarium*, but higher than *GAPDH Ks* between *V. dahliae* and *V. alfalfae* (Figure S13B), indicating that the cross-order HGT occurred prior to *Verticillium* species diversification. Additionally, lower *K_s_* values between *D* orthologs of *V. dahliae* and *V. alfalfae* relative to *GAPDH* suggest a more recent cross-species HGT within *Verticillium*. Together, these results indicate multiple horizontal transfers of *D*, including between *Verticillium* and *Fusarium* and among *Verticillium* species.

*Starships* are large DNA transposons carrying tens to hundreds of cargo genes and are widespread in Pezizomycotina fungi^35–37^. They have been implicated in HGT between fungal genera, including virulence genes between *Verticillium* and *Fusarium*^38,39^. To assess whether *D* homologs were transferred via *Starships*, previously identified *Starships* in *Verticillium* and *Fusarium* were analyzed^38,40^. In *Verticillium*, the *D* gene is located on *Ar1h2* haplotype *Starships*, whereas *D-like* orthologs are not (Figure 6A). *Ar1h2 Starships* (∼0.5 Mb) contain five tyrosine recombinase “captain” genes mediating *Starship* transposition, suggesting nested insertions. Notably, 74% of *Ar1h2* regions share >98% nucleotide identity with 64% of *Ar4h1 Starships*, which carry the gene encoding the horizontally transferred Av2 effector^38^, indicating divergence from a common ancestor. Furthermore, *Ar1h2 Starship* regions are >99% identical between *V. dahliae* and *V. alfalfae*, despite a genome-wide average identity of 93%, suggesting recent horizontal transfer between species. These findings implicate *Ar1h2 Starships* in the horizontal transfer of the *D* gene among *Verticillium* species.

**Fig. 6.**
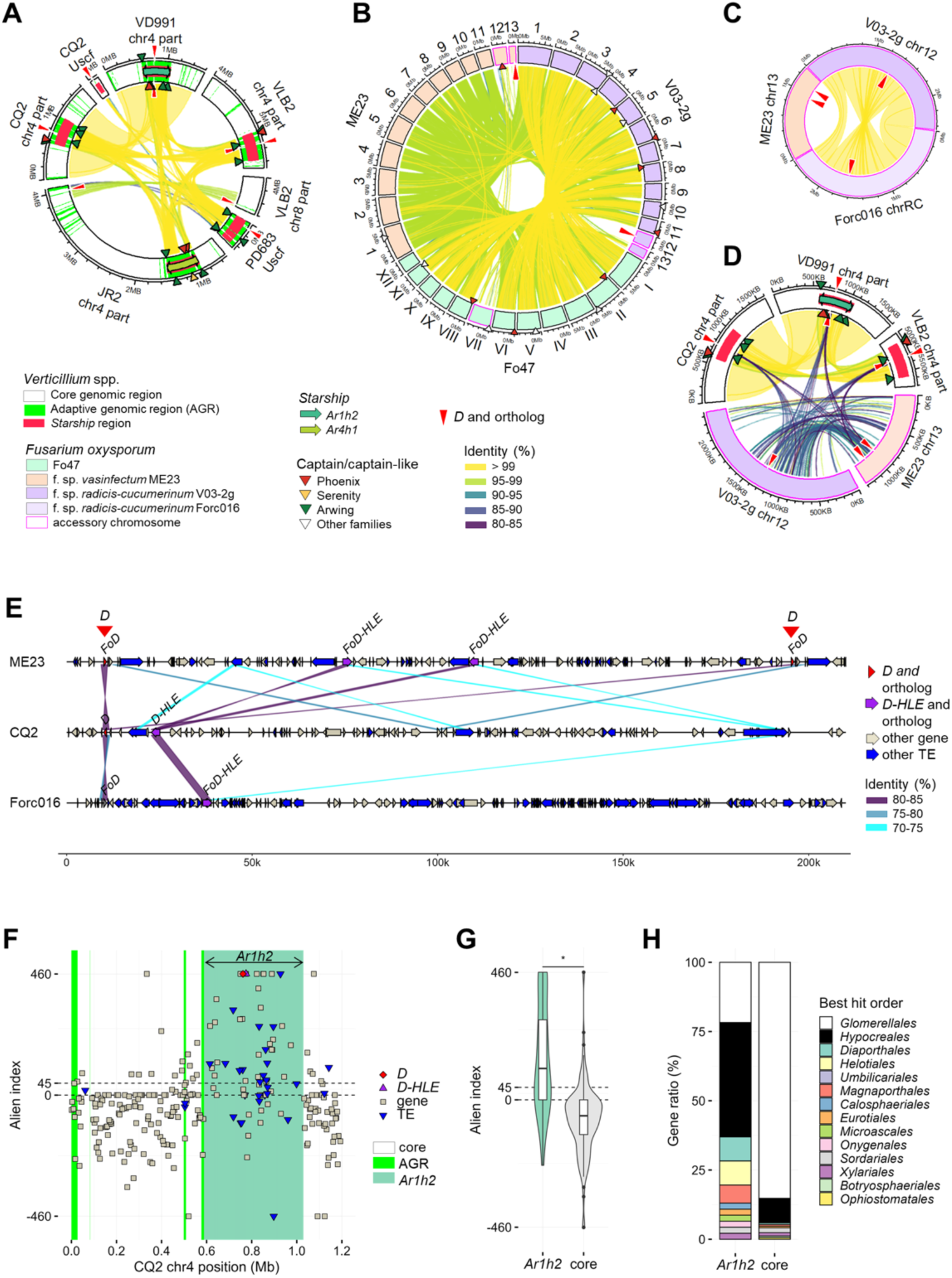
*Starship*-mediated *D* gene transfer and its association with a *Helitron*-like transposon. A-D, Circular plots showing synteny among genomic regions of *Verticillium* and *Fusarium* strains. Red elongated arrowheads indicate positions of *D* homologs sharing >89% nucleotide sequence identity over 99% coverage. Tracks in *Verticillium* genomic regions are colored to indicate conserved core genomic regions or plastic adaptive genomic regions (AGRs) and are overlaid with bold lines and arrows representing *Starship* syntenic regions and *Starships*, with *Starships* colored by haplotype. AGRs exhibit chromatin profiles similar to those of *Starship* regions, and some AGRs may represent remnants of ancestral *Starships* (Cook et al., 2020; Sato et al., 2025). Regular triangles indicate captain and captain-like genes, colored by family. Ribbons connect homologous regions sharing >80% nucleotide sequence identity over 10 kb in (A-C) or over 1 kb in (D). (A) Synteny among *D*-carrying genomic regions in *Verticillium dahliae* strains CQ2, VD991, and JR2, *V. longisporum* strain VLB2, and *V. alfalfae* strain PD683. (B) Synteny among complete genome assemblies of *Fusarium oxysporum* strains Fo47, ME23, and V03-2g. (C) Synteny among *D*-carrying accessory chromosomes in *F. oxysporum*strains ME23, V03-2g, and Forc016. (D) Synteny among *D*-carrying genomic regions in *Verticillium* spp. and accessory chromosomes in *F. oxysporum*. (E) Location of *D* and *D-HLE* (*D-proximal Helitron-like Element*) in *V. dahliae* strain CQ2 and their orthologs (*FoD* and *FoD-HLE*) in *F. oxysporum* strains ME23 and Forc016. Ribbons connect homologous regions sharing >70% nucleotide sequence identity over 0.5 kb. (F) Genomic positions and inferred potential for cross-order horizontal gene transfer (HGT) of genes and transposable elements (TEs) surrounding the *D*-carrying *Ar1h2 Starship* in *V. dahliae* CQ2. Alien index (AI) values >45 suggest HGT, whereas values <0 suggest vertical inheritance. (G) Violin and box plots summarizing AI values of genes within the *Ar1h2 Starship* and flanking core genomic regions shown in (F). Black points indicate outliers. Center, upper, and lower lines in box plots denote the median, third quartile, and first quartile, respectively. Asterisks indicate a significant difference (two-sided Brunner-Munzel test, *p* < 2.2e-16, *n* = 46 in *Ar1h2* and *n* = 169 in core regions). (H) Stacked bar plots showing the percentage of *Verticillium* genes whose closest orthologs are found in genomes from the indicated orders of Pezizomycotina, excluding *Verticillium*, within the order Glomerellales. The query genes are the same as those analyzed in (G).

In *Fusarium*, no *D* orthologs were found within previously identified *Starships*, nor were intact *Starships* or captain-like genes detected on chromosomes or scaffolds carrying *D*. However, in complete telomere-to-telomere assemblies of *F. oxysporum* f. sp. *vasinfectum* and f. sp. *radicis-cucumerinum*^41–43^, *D* orthologs reside on distinct accessory chromosomes (Figure 6B), one of which closely resembles an accessory chromosome previously shown to transfer horizontally between *F. oxysporum* strains and confer pathogenicity^44^ (Figure 6C). Direct evidence of horizontal *Starship* transfer is difficult to obtain for ancient events, as *Starship* regions are highly plastic and may become unrecognizable due to rearrangements and active transposable elements (TEs)^38,45^. We therefore assessed whether *Ar1h2 Starships* share similarity with *Fusarium* regions surrounding *D*. While large-scale synteny is absent, partial sequence similarity exists at several loci, notably at the *D* locus and a nearby *Helitron*-like TE (*D-HLE*) ∼12 kb from *D*, with intervening regions populated by non-syntenic TEs (Figure 6D–E). Helitrons transpose via rolling-circle replication and can capture adjacent sequences^46–48^. *D-HLE* is found exclusively in *Verticillium* and multiple Hypocreales genera, including *Fusarium*, and exhibits an AI score of 460.5, comparable to *D* itself, suggesting horizontal transfer (Figure S14). Thus, *D-HLE* could have mobilized *D* between an *Ar1h2 Starship* and an ancestral accessory chromosome.

To assess whether additional *Ar1h2* cargo genes were co-transferred, cross-order AI scores were calculated for all *Ar1h2* cargo elements and compared with flanking core genes. The median AI score of cargo genes (113) far exceeded that of flanking genes (−58), indicating horizontal transfer of most *Ar1h2* cargo genes (Figure 6F–G). Among Pezizomycotina, 41% of the most conserved *Ar1h2* cargo orthologs are found in Hypocreales, including *Fusarium*, spanning nine additional genera (Figure 6H, S14). Currently, these cargo genes are dispersed across chromosomes/scaffolds with frequent presence/absence variation (Table S10), consistent with post-transfer *Starship* degradation. Collectively, these findings support a model in which the entire *Ar1h2 Starship*, rather than *D* and *D-HLE* alone, was horizontally transferred between *Verticillium* and Hypocreales, followed by *D-HLE*–mediated transposition of *D* and extensive post-transfer rearrangements that disrupted synteny at *D* loci between the genera.

### *Fusarium* D homologs induce wilting on cotton and cucurbit plants

To test whether *Fusarium* D effector homologs can induce cotton defoliation, proteins from the cotton-pathogenic *F. oxysporum* f. sp. *vasinfectum* (*DFov*, strain FovNRRL31665) and the cucurbit-pathogenic *F. oxysporum* f. sp. *radicis-cucumerinum* (*DForc*, strain Forc016, which cannot infect cotton) were produced and applied to cotton seedlings in MS medium at 5 μM, with *V. dahliae* D protein and BSA as positive and negative controls, respectively. Both *DFov* and *DForc* induced wilting within two days and cotyledon detachment after seven days, similar to the *V. dahliae* D protein, while BSA had no effect (Figure 7). This demonstrates that *DFov* and *DForc* encode functional effectors capable of triggering cotton defoliation, even when produced by a fungus unable to infect cotton.

**Figure 7.**
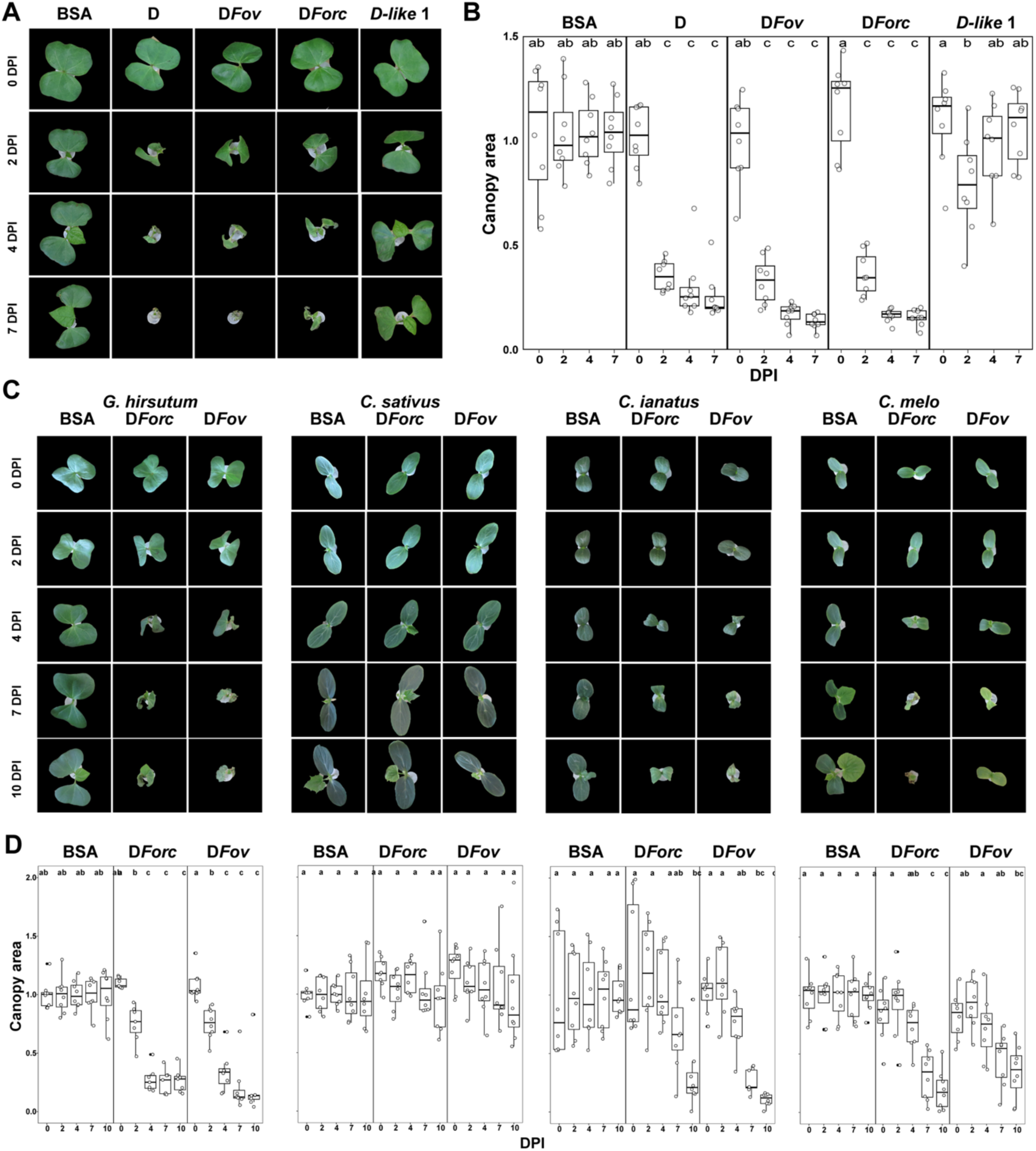
D homologs of *Fusarium oxysporum* differently affect cucurbit species. (A) Typical appearance of cotton seedlings (cv. Xinluzao63) upon application of D effector protein homologs from *Verticillium dahliae* (*D*, *D-like1*; 5 µM), *Fusarium oxysporum* f. sp. *vasinfectum* (D*Fov*; 5 µM), *F. oxysporum* f. sp. *radicis-cucumerinum* (D*Forc*; 5 µM) at 0, 2, 4 and 7 days post treatment. Bovine serum albumin (BSA; 5 µM) was used as control. (B) Quantification of the canopy area of treatments shown in panel (A) normalised by the average of BSA treated plants at 0, 2, 4 and 7 days after start of the treatment. Each bar represents eight plants. Different letter labels indicate statistically significant differences (one-way ANOVA and Tukey test). Experiments were repeated twice with similar results. (C) Typical appearance of *G. hirsutum* (cotton, cv. Xinluzao63), *C. sativus* (cucumber, cv. Paraiso), *C. ianatus* (watermelon, cv. Black diamond) and *C. melo* (melon, cv. Brotmabon), upon application of D effector protein homologs from *Fusarium oxysporum* f. sp. *radicis-cucumerinum* (D*Forc*; 5 μM), *F. oxysporum* f. sp. *vasinfectum* (D*Fov*; 5 μM) at 0, 2, 4,7 and 10 days post treatment. Bovine serum albumin (BSA; 5 μM) was used as control. (D) Quantification of the canopy area of treatments shown in panel (C) normalised by the average of BSA treated plants at 0, 2, 4, 7 and 10 days after start of the treatment. Each bar represents eight plants. Different letter labels indicate statistically significant differences (one-way ANOVA and Tukey test). Experiments were repeated twice with similar results.

The effect of DFov and DForc was also tested on cucurbit hosts (cucumber, watermelon, and melon). Both proteins caused wilting in watermelon and melon within two days, followed by leaf shrinkage and curling after seven days, whereas cucumber plants remained unaffected (Figure 7C–D). These results confirm that *Fusarium* D homologs are functional effectors capable of inducing host-specific disease symptoms despite sequence differences.

### The D effector displays a unique fold

To investigate the *in planta* function of D proteins, structural predictions for all functional D homologs were attempted using AlphaFold3^49^, but only low-confidence models were obtained (pLDDT < 50). Consequently, experimental structure determination was pursued via vapor-diffusion crystallization. Circular dichroism (CD) spectroscopy indicated that the D protein is predominantly composed of β-sheets (Figure S15A), while size-exclusion chromatography (SEC) showed it exists as a monomer (Figure S15B). Crystallization trials of D homologs from *V. dahliae*, *F. oxysporum* f. sp. *vasinfectum* (D*Fov*), and *F. oxysporum* f. sp. *radicis-cucumerinum* yielded diffraction-quality crystals only for DFov.

The D*Fov* structure was solved by X-ray crystallography at 1.5 Å resolution using single-wavelength anomalous diffraction (SAD) procedure with Hg atoms. The protein crystallized in space group P2₁ with one molecule per asymmetric unit and 33% solvent (Table S9). Residues 1–18, 43–50, and 100–107 were unstructured and omitted. D*Fov* forms an elongated monomeric β-barrel composed of eleven β-strands organized into three antiparallel β-sheets (B1: β1β2β10, B2: β3β4β5β11, B3: β6β7β8β9) and three α-helices. B2 and B3 pack to form the β-barrel, B1 and α3 cap one end, while α1 and α2 connect β2 to β3 on the opposite side (Figure 8A–B). The β-barrel is solvent-inaccessible, with a predominantly negative surface and a cluster of positive residues on one side (Figure 8C).

**Figure 8.**
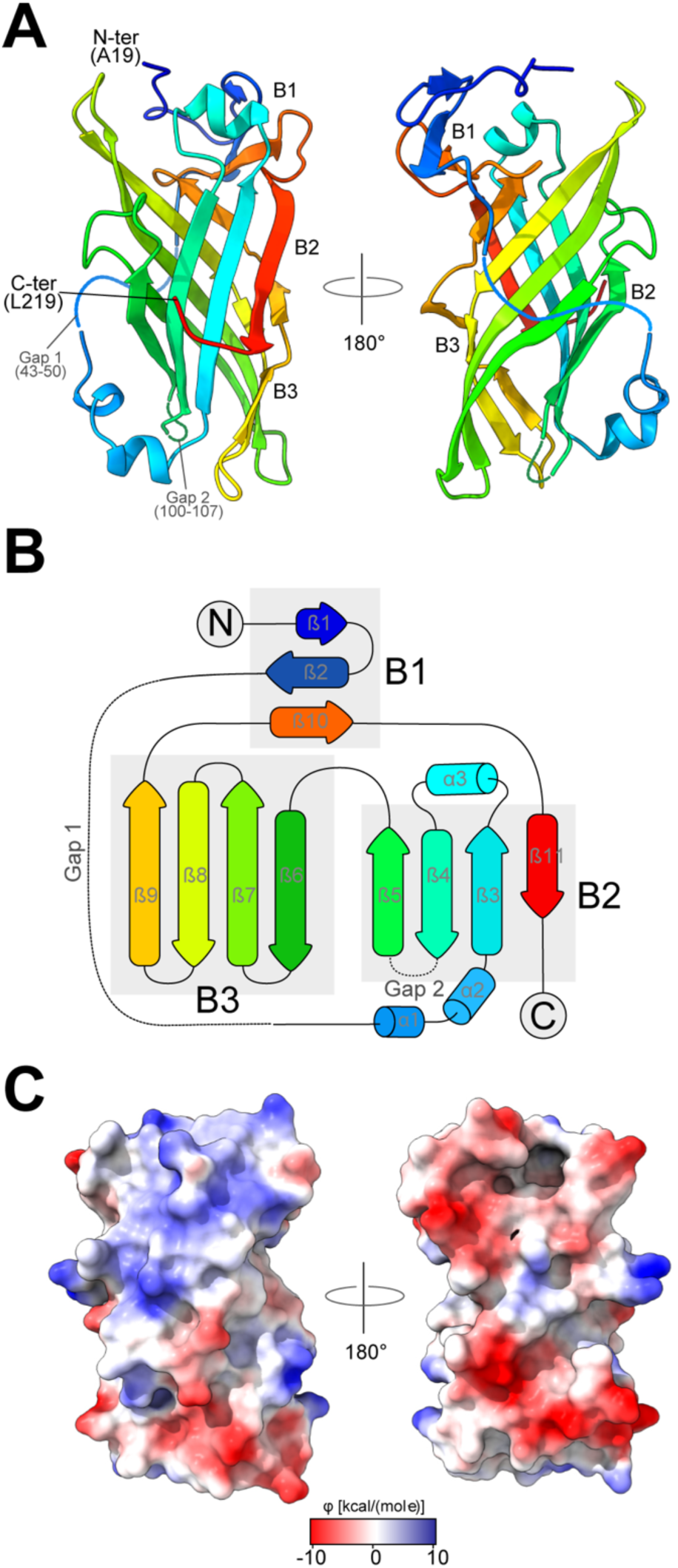
Crystal structure of the *F. oxysporum* f. sp. *vasinfectum* D protein homolog. (A) Cartoon representation of D*Fov* crystal structure coloured according to the standard rainbow mode, with blue and red indicating the N-terminal and C-terminal regions, respectively. The first 19 N-terminal residues and 2 internal regions (indicated as dotted lines and named Gap 1 and 2) are not visible due to a high conformational mobility. (B) Fold topology of the different secondary structure elements of the D*Fov* fold. The three β-sheets are indicated as B1-3. B2 and B3 interact to form the β-barrel core of the protein. (C) Electrostatic surface potential of D*Fov* is calculated from atomic partial charges and coordinates according to Coulomb’s law: φ = Σ [q_i_ / (εd_i_)] using a default dielectric constant of 4.0 and a distance of 1.4 Å. Positive and negative regions are coloured in blue and red, respectively. The surface of D*Fov* is anisotropically charged with a prevalence of positive or negative charge on opposite sides.

Comparison with the low-confidence AlphaFold3 model using TM-align revealed poor structural similarity (TM-score < 0.5), confirming the model’s low accuracy (Figure S16). Structural homology searches via the Dali server^50^ identified a distant similarity to the *Parastagonospora nodorum* effector Tox3 (PDB: 6WES, Z-score 5.3; Outram et al., 2021). To expand the search, D*Fov* was compared against a database of ∼5,000 AlphaFold2-predicted fungal effector structures ^52^, revealing high-confidence structural homologs (Z-scores up to 12.4) only in *Claviceps purpurea*, *Epichloe festucae*, and *F. oxysporum* f. sp. *conglutinans* (Table S11). The structural similarity network placed D*Fov* in a cluster with these predicted effectors, indicating that D*Fov* adopts a fold conserved across diverse fungal lineages but dissimilar to previously determined effector protein structures (Figure S17). These results provide the first experimental evidence for this structural class, validating its existence beyond *in silico* predictions and establishing a framework for functional characterization of D effector proteins.

## DISCUSSION

The pathogenicity of plant-pathogenic fungi is mediated by secreted effector proteins that manipulate host physiology to promote infection^16,17,53–55^, although more recent evidence indicates that effectors can also target host-associated microbiota to promote infection^22^. Effector functions are often redundant, such that loss of individual effectors is typically dispensable for disease. Nevertheless, several studies demonstrate that a single effector can determine pathogenicity on specific hosts. For instance, the transcriptional activator-like effector PthA from *Xanthomonas citri* confers citrus pathogenicity when transferred into non-pathogenic strains^56–58^, and the necrotrophic effector SnTox1 enables *Parastagonospora nodorum* to infect wheat lines carrying the susceptibility gene *Snn1*^26,59^.

Although *Verticillium dahliae* strains collectively display a broad host range, pathogenicity and symptom severity vary widely^60^. Defoliating (D) and non-defoliating (ND) pathotypes have long been described in cotton– and olive-infecting strains^5,6^, yet the genetic basis of defoliation remained unclear. Although defoliation has been proposed to involve a horizontally transferred gene cluster from *Fusarium oxysporum* f. sp. *vasinfectum*, including secondary metabolite–associated genes, no single causal gene had been identified^61^. Here, we show that a single effector, termed the D effector, governs pathogenicity and defoliation in cotton and olive. Deletion of both *D* gene copies abolished defoliation, introduction into an ND strain conferred this trait. Moreover, importantly, application of the heterologously produced effector protein alone induced defoliation, demonstrating that the D effector directly drives symptom development.

*Verticillium dahliae* is regarded as an asexual species because no sexual cycle has been observed and mating type distribution is highly skewed, resulting in a predominantly clonal population structure^3,62–66^. This clonality was originally described using vegetative compatibility groups (VCGs)^67,68^, with isolates in clonal lineage VCG1A corresponding to the defoliating (D) pathotype, while non-defoliating (ND) strains occur across all other lineages^8,32,69,70^. The D pathotype is thought to have originated in North America before dispersing globally^71^. Consistent with this, SNP-based genotyping of a worldwide collection of *V. dahliae* isolates from diverse hosts showed that D pathotype strains from Europe, North America, and China all cluster within VCG1A and share nearly identical SNP haplotypes, supporting a single origin followed by intercontinental spread^11,65^.

Plant pathogens continuously reshape their effector repertoires to evade host immunity and sustain aggressiveness^16,55^. The emergence of new effector genes can enhance virulence or expand host range^72,73^ and is driven by mechanisms such as genome hybridization^74^, gene duplication^75^, and horizontal gene transfer^72,76^. *Verticillium dahliae* exhibits a two-speed genome in which segmental duplications generate adaptive regions enriched in *in planta*–expressed effector genes, implicating duplication as a major force in virulence evolution^77,78^. In this study, we show that the *D* effector gene arose by a recent segmental duplication, yielding two identical copies with highly similar flanking sequences. Quantitative phenotyping demonstrated that each copy contributes to fungal aggressiveness, indicating that duplication of the D effector is important for maintaining high virulence in D pathotype strains. More broadly, duplicated effector genes in filamentous pathogens often follow a duplication–divergence trajectory, in which redundancy allows one copy to evolve new functions, as observed for Crinkler effectors in *Phytophthora sojae*^79^ and effector diversification in *Ustilago maydis*^75^. Such processes can ultimately drive extensive effector expansions, exemplified by *Blumeria graminis*, where many effectors share an RNase-like fold derived from a single ancestral gene^80,81^.

A key finding of this study is the identification of functional *D* effector homologs in multiple *formae speciales* of *Fusarium oxysporum*. Although *V. dahliae* is a broad host-range pathogen and *F. oxysporum* strains are host specialists, both share a similar infection strategy as soil-borne vascular pathogens and possess overlapping effector repertoires, including Ave1 and Av2^21,76,82^. Recent genomic analyses have implicated giant *Starship* transposons in the horizontal transfer of these virulence effectors^38^, and the D effector identified here represents an additional effector shared between the two genera. Our evolutionary analyses highlight *Starships* as major vectors of virulence gene transfer and reveal a previously unrecognized evolutionary link between *Starships* and pathogenicity-associated accessory chromosomes in *Fusarium*. The presence of nearly identical *D*-carrying *Starships* across *Verticillium* species supports recent *Starship*-mediated transfer within the genus, whereas sequence divergence patterns and *K_s_* estimates indicate a more ancient transfer to or from *Fusarium*, where intact *Starships* have degenerated, but their cargo persists. In *Fusarium*, *D* orthologs reside on accessory chromosomes that are enriched in virulence genes and capable of horizontal chromosome transfer^83,84^. Similar patterns are observed for other *Starship* cargo, such as *spore killing* (*spok*) genes^85^, with *Starship* degradation attributed to genomic rearrangements, loss of captain genes, and repeat-induced point (RIP) mutation activity^36,37,45^. *Fusarium* accessory chromosomes are enriched in rearrangements, segmental duplications, and in transposon activity that may have contributed to the accelerated degeneration of *Starships*^86,87^. Our data further suggest that Helitron-like transposable elements facilitated movement of *Starship* cargo to and from accessory chromosomes, paralleling TE-mediated mobilization of effectors such as *ToxA* and *ToxB* in other fungi^88–90^. Together, these findings support a model in which coordinated interactions among *Starships*, transposable elements, and accessory chromosomes drive the spread and diversification of fungal virulence genes.

Despite advances in structural prediction, including AlphaFold3, no reliable model could be generated for the D effector. Determination of the 3D crystal structure of its *Fusarium* homolog (D*Fov*) revealed a previously uncharacterized protein fold with a weak structural homology to SnTOX3 in the Protein Data Bank, placing the D effector in a distinct structural class and raising questions about its mechanism of action. Its activity across diverse hosts—cotton, olive, *Nicotiana benthamiana*, and *Arabidopsis thaliana*—suggests it targets a conserved plant component involved in core immune or developmental processes. Interestingly, the D effector, also identified as the “TRADE” effector in *V. longisporum*, triggers bundle sheath cell identity switches into tracheary elements in *A. thaliana*, enhancing water storage and drought resilience, likely via interaction with VARICOSE (VCS), a conserved mRNA turnover component^91^. Whether this mechanism underlies cotton and olive defoliation remains to be confirmed.

Despite the severe impact of D pathotype strains and the occurrence of *D-like* genes in all thus far characterised *V. dahliae* strains, no resistance (*R*) genes have been identified in cotton or olive^92,93^. Because effectors can reveal host recognition, they are widely used as probes to identify *R* genes through hypersensitive responses in germplasm screening^94–97^. Accordingly, the D effector represents a promising tool for discovering *R* gene–mediated resistance in cotton, olive, and other crops.

## MATERIALS AND METHODS

### Genome sequencing and deep transcriptome sequencing

The genome of *V. dahliae* strain CQ2 (Table S1) was sequenced using PacBio SMRT long-read technology, with library construction as previously described^27,28,33^. *V. dahliae* strains Vd39, V4, BP2, V574, V700, V117, V76, ST.100, T9, and V991 (Table S1) were sequenced on the Illumina HiSeq 2000 following library preparation with 500-bp inserts and 100-bp paired-end reads at the Beijing Genome Institute (BGI, China).

Phylogenetic relationships among *V. dahliae* strains were inferred using REALPHY v1.12^98^, with genomic reads mapped to the gapless reference genome of strain JR2^33^ using Bowtie2^99^. A maximum-likelihood tree was constructed with RAxML v8.2.8 under the GTRGAMMA model and 500 bootstrap replicates^100^, and rooted with *V. alfalfae* Ms102.

For transcriptome analysis, 12-day-old cotton seedlings were root-inoculated with conidiospores of the D pathotype strain V991 as previously described^101^. Stems were harvested at 6, 9, 12, and 15 days post inoculation for total RNA extraction using the Spectrum Plant Total RNA Kit (Sigma-Aldrich, USA). cDNA synthesis, library preparation with 200-bp inserts, and Illumina sequencing (90-bp paired-end reads) were performed at BGI (Shenzhen, China).

### *V. dahliae* comparative genomics

The *V. dahliae* strain CQ2 genome was assembled with HGAP v3.0 using default parameters^102^ and annotated with MAKER2 v2^103^, followed by manual curation of gene models in regions of interest, including verification of MAKER2 predictions and refinement of exon–intron boundaries using transcriptome data. To identify genes associated with cotton defoliation, Illumina reads from defoliating (D) and non-defoliating (ND) pathotype strains (Table S1), together with sequence data from 72 *V. dahliae* strains from NCBI (project PRJNA171348), were mapped to the CQ2 genome using BWA with default settings^104^. Presence/absence analysis was performed with BEDtools^105^ and R^106^ by partitioning the genome into 100-bp bins and assessing read presence in 50-bp sliding windows, identifying regions covered by D strains but lacking ND strain coverage. Coverage was normalized as (reads per bin × 10,000,000) / total mapped reads, evaluated per sample, and filtered using a cutoff of 5. Genes located in lineage-specific (LS) regions unique to D pathotype strains were extracted using BEDtools intersect v2.25^105^, and bioinformatic predictions in these LS regions were validated by mapping RNA-seq reads from cotton plants infected with D pathotype strain V991 using TopHat v1.4.0^107^.

### Gene expression analysis

To assess *D* gene expression in *D* deletion and complementation strains, *V. dahliae* cultures were grown in liquid Czapek–Dox medium at 22 °C for one week with shaking at 150 rpm, after which mycelia and conidia were harvested for RNA extraction. First-strand cDNA was synthesized using the MMLV reverse transcriptase system (Promega, USA). RT-PCR was performed with primers CQ2D-F and CQ2D-R (Table S2) in 25 μl reactions containing 17.9 μl sterile water, 5 μl 5× PCR buffer, 0.5 μl dNTPs, 0.5 μl of each primer, 0.1 μl GoTaq DNA polymerase (Promega, USA), and 1.0 μl first-strand cDNA (100 ng μl⁻¹). The *V. dahliae GAPDH* gene served as an endogenous loading control, and PCR products were analyzed by agarose gel electrophoresis.

### BLAST searches and protein sequence alignment

The genomes of 63 sequenced *V. dahliae* strains (Table S1) were screened for *D*-*like* alleles using BLASTn with default parameters^108^. Matching sequences were extracted with the BEDtools getfasta utility^105^ and aligned using webPRANK^109^ to assess allelic variation. Predicted amino acid sequences encoded by the *D* gene and its *D*-*like* alleles were also aligned with webPRANK^109^ and visualized using ESPript v3.0^110^.

### Heterologous D effector protein production and cotton bioassays

The D effector protein was produced in *Pichia pastoris* as previously described(Sánchez-Vallet et al., 2013; Kombrink et al., 2017b). The *D* gene was PCR-amplified with primers adding N-terminal His and FLAG tags (Table S2) and cloned into the *P. pastoris* expression vector pPIC9 (Invitrogen, Carlsbad, CA, USA), which was transformed into strain GS115. Putative transformants were screened for D protein expression in BMM medium, and one clone was selected for large-scale cultivation in a BioFlo 120 fermenter (Eppendorf AG, Hamburg, Germany). The His-tagged D protein was purified using a Ni²⁺-NTA Superflow column (Qiagen, Venlo, the Netherlands), dialyzed against 100 mM NaCl, quantified by absorbance at 280 nm, and analyzed by SDS–PAGE and Western blotting with a mouse anti-His monoclonal antibody.

To evaluate the effect of the D effector, three-week-old cotton seedlings were uprooted, rinsed, and their roots immersed in 10 mL of D protein solution (8.96 μM). Water and the *V. dahliae* chitin-binding effector Vd2LysM (8.96 μM) were used as negative controls. Seedlings were incubated in a growth chamber at 21/19 °C (16/8 h day/night), 70% relative humidity, for 16 days and monitored for symptom development.

### D homolog identification

Genomes of approximately 22,000 fungi were downloaded from NCBI, and the *D* gene sequence from *V. dahliae* strain CQ2 was used to query these genomes with Exonerate v2.3.0^112^ using the protein2genome model and a 50% identity threshold. Exonerate output gff files were subsequently processed with gffread v0.11.7^113^ together with the corresponding genome sequences to extract predicted protein sequences (Table S3).

### Phylogenetic analysis

Nucleotide sequences of *D* homologs were extracted from genome assemblies listed in Table S3 using SeqKit v2.10.0 ^114^, with assemblies downloaded from NCBI via NCBI Datasets v18.6.0 ^115^. Sequences were aligned using the L-INS-i algorithm in MAFFT v7.526 with the *adjustdirection* option ^116^, and ambiguously aligned sites were removed with TrimAl v1.5.rev0 using the *– automated1* option, retaining 740 aligned residues ^117^. Phylogenetic inference was performed with IQ-TREE v3.0.1 ^118^, which automatically selected the best-fit nucleotide substitution model (TPM2u+F+G4) using ModelFinder ^119^, and branch support was estimated with 1,000 ultrafast bootstrap replicates ^120^. The consensus tree was visualized using the R package *ggtree* v3.14.0 ^121^, and the closest *Fusarium* ortholog to D was identified using the *cophenetic.phylo* function in the R package *ape* v5.8.1 ^122^. Synonymous substitution rates (Ks) were calculated with KaKs_Calculator v2.0 using the MA model ^123^ on coding sequences of D and GAPDH homologs extracted from coordinates listed in Table S4.

### Detection and visualization of genomic syntenies

Accession numbers of genome assemblies used for synteny detection are listed in Table S7. Genes and TEs of *V. dahliae* strain CQ2 were annotated as mentioned above. Genes of *F. oxysporum* strains ME23 and Forc16 were annotated in previous studies^41,43,124^, while TEs in these strains were annotated in this study using EDTA v2.2.1 with the options ‘––overwrite 1 ––sensitive 1 ––anno 1’ ^125^. *Verticillium* genomic regions (core regions, AGRs, and *Starship* regions) and synteny between *Verticillium* genomes were identified previously^38^. Synteny between *Fusarium* genomes and between *Verticillium* and *Fusarium* genomes was detected by pairwise genome sequence alignment using ‘nucmer’ with the ‘––maxmatch’ option, followed by alignment filtering using delta-filter with the ‘-m’ option and coordinate extraction using ‘show-coords’ with the ‘-THrcl’ option in MUMmer v4.0.1^126^. The genomic synteny was visualized with R package ‘circlize’ v0.4.16^127^. Local sequence similarity was detected by pairwise genome sequence alignment using BLASTn with the same options mentioned above and visualized together with gene and TE loci using R package ‘gggenomes’ v1.0.1^128^.

### Identification of *Starships* and captain genes

*Starships* and captain-like genes in *Verticillium* shown in the figures were identified previously^38^. *Starships* reported from diverse Pezizomycotina fungi^38,40^ were screened for *D* homologs using BLASTn with the parameters described above, which identified no *Fusarium Starships* carrying *D*. Because additional genome assemblies have since become available, all currently available *Fusarium* assemblies in NCBI that contain *D* homologs (Table S3) were further examined for *Starships* and captain-like genes. *Starship* detection was performed using the ‘starfish*’* v1.1.0 pipeline with previously described settings, which identifies insertion–deletion polymorphisms associated with 5′-end captain genes^40^. Captain-like genes were detected using a Pezizomycotina captain and captain-like protein database via translated nucleotide similarity searches implemented in MetaEuk v6.a5d39d9^129^. No captain candidates were identified on *D*-containing chromosomes or scaffolds (Table S8), indicating the absence of detectable *Starships*.

### Disease assays

Cotton (*Gossypium hirsutum* cv. Xinluzao63), *Nicotiana benthamiana*, and *Arabidopsis thaliana* (Col-0) were grown under controlled greenhouse conditions (Unifarm, Wageningen, the Netherlands). Olive (*Olea europaea* cv. Picual) was grown in a greenhouse under natural lighting and day/night temperature of 27/21°C (Córdoba, Spain). Two-week-old cotton and *A. thaliana*, three-week-old *N. benthamiana* and eight-month-old olive plants were used for inoculation. *Verticillium dahliae* strains (Table S1) were grown on potato dextrose agar (PDA) at 22°C for 7-10 days. Conidiospores were collected from PDA and washed with tap water for inoculation. Disease assays on cotton, *N. benthamiana* and *A. thaliana* were performed using the root-dip method as previously described ^130,131^. Olive assays followed published protocols^132,133^. Symptoms were scored up to 21 (*A. thaliana* and *N. benthamiana*), 28 (cotton), or 132 (olive) days post inoculation (dpi). *A. thaliana* and *N. benthamiana* were photographed, and canopy area (*N. benthamiana*) and total rosette area (*A. thaliana*) were quantified with ImageJ. Cotton height was measured with a rectilinear scale, and defoliation was classified as 0 (0%), 1 (<25%), 2 (<50%), 3 (<75%), or 4 (<100%) leaf drop^26,134^. Fungal biomass in *N. benthamiana*, *A. thaliana*, and cotton was quantified by real-time PCR^130,131^. Olive disease severity was assessed twice weekly based on the percentage of leaves with chlorosis or defoliation: 0 (no symptoms), 1 (1–33%), 2 (34–66%), 3 (67–100%), 4 (dead plant)^31,133^. Disease incidence (DI), mortality (M), and disease intensity index (DII) were calculated per treatment. DII was defined as DII = (ΣSi × Ni)/(4 × Nt), where Si = symptom severity, Ni = number of plants with severity Si, and Nt = total plants. Final DI was the percentage of affected plants at the end of the bioassay. The area under the disease progress curve (AUDPC) of DII over time^135^ and final severity were calculated. Data were analyzed by ANOVA, and means were compared using Fisher’s protected LSD (P = 0.05) in Statistix v10.0 (Analytical Software, 1985–2013).

### Generation of *D* gene deletion strains

To generate a *D* single gene deletion construct, sequence stretches of approximately 1.2 kb upstream and 1.3 kb downstream of the D coding sequence were amplified from genomic DNA of D pathotype strain CQ2, using primer pairs SKO-D-LBF/LBR and SKO-D-RBF/RBR (Table S2). The amplicons were cloned into vector pRF-HU2 as described previously^136^, and the resulting deletion construct was transformed into CQ2 via *Agrobacterium tumefaciens*-mediated transformation^137^. Putative deletion transformants were selected on PDA containing 200 μg/mL cefotaxime and 50 μg/mL hygromycin B (Duchefa, Haarlem, The Netherlands) and homologous gene replacement was verified with PCR analysis using outside primer-F and outside primer-R (Table S2). To generate *D* double gene deletion mutants, sequence stretches of approximately 1.1

kb upstream and 1.2 kb downstream of the *D* coding sequence were amplified using primer pairs DKO-D-LBF/LBR and DKO-D-RBF/RBR (Table S2). The amplified products were cloned into vector pRF-NU2. Next, the gene replacement construct was transformed into a *D* gene single deletion mutant. Putative double deletion transformants were selected on PDA containing 50 μg/mL hygromycin B and 15 μg/mL nourseothricin (Sigma-Aldrich Chemie GmbH, Buchs, Switzerland), and were subjected to PCR to confirm genuine double *D* gene deletions using outside primer-F and outside primer-R (Table S2). Reverse transcription-PCR (RT-PCR) was used to confirm no *D* gene transcripts in *D* double deletion mutants using primers D-RT-F and D-RT-R, and *V. dahliae* GAPDH (glyceraldehyde-3-phosphate dehydrogenase) gene as an endogenous control (Table S2). Meanwhile, the same *D* single deletion construct and double deletion constructs were used to generate *D* single and *D* double deletion mutants in *V. dahliae* strain V150I, which causes defoliation on olive^32^. *D* gene complementation was performed by using a genomic construct consisting of the *D* gene coding sequence with 1.1 kb upstream and 1.2 kb downstream sequences (pD::D) using primer pairs D-com-F and D-com-R (Table S2). The amplified products were cloned into GatewayTM compatible vector PCG using a standard BP reaction. The gene complementation construct was further transformed into a *D* single deletion mutant, a *D* double deletion mutant, a ND pathotype strain JR2, respectively. Putative *D* gene complementation transformants were selected on PDA supplemented with 200 μg/mL of cefotaxime and 25 μg/mL geneticin (Sigma-Aldrich Chemie BV, Zwijndrecht, The Netherlands). Reverse transcription-PCR (RT-PCR) was used to examine *D* gene transcription in these transformants.

### Heterologous production of D homologs in *Escherichia coli*

The nucleotide sequences encoding mature D protein from *Verticillium dahliae* strain CQ2, D-like 1 from *V. dahliae* strain JR2 and D homologs from *F. oxysporum* f. sp. *vasinfectum* strain *Fov*NRRL31665 (D*Fov*) and from *F. oxysporum* f. sp. *radicis-cucumerinum* strain *Forc*016 (D*Forc*) without predicted signal peptides were synthesised by Eurofins Genomics (Louisville, KY, USA) and amplified with PCR using primers listed in (Table S2). The resulting amplicons were cloned into the expression vector pET-28a (Novagen, Madison, USA), sequenced, and transformed into chemo-competent *E. coli* strain BL21 cells. Single colonies were picked from selection medium and grown overnight in LB (10 g/L tryptone, 5 g/L yeast extract, 5 g/L NaCl) supplemented with 50 µg/mL kanamycin at 37°C and 200 rpm. The optical density (OD_600_) of the overnight cultures was determined spectrophotometrically (BioPhotometer, Eppendorf, Hamburg,

Germany). Next, 1 L cultures in LB medium were initiated at an OD_600_ of 0.1 by adding an appropriate volume of overnight culture and subsequently incubated at 37°C and 200 rpm until an OD_600_ of ∼0.6 was reached. Then, protein expression was induced by addition of 100 µL of a 1 M aqueous solution of IPTG (Duchefa, Haarlem, the Netherlands) to reach a final concentration of 100 µM, and cultures were incubated overnight at 37°C and 200 rpm. Subsequently, cultures were pelleted for 30 minutes at 5,000 x g at room temperature. Next, pelleted cells were resuspended in lysis buffer (50 mM Tris, 200 mM NaCl, 10% [v:v] glycerol, 6 mg/mL lysozyme, 2 mg/mL sodium deoxycholate, 20 μg/mL DNase, 1 mM PMSF; pH 8.0) and lysed for one hour on ice. The lysate was then centrifuged for 30 minutes at 21,000 x g at 4°C. The supernatant was transferred to a new tube and subsequently loaded onto an affinity purification column (His60 Ni Superflow, TaKaRa, Gotenburg, Sweden) that was first equilibrated with equilibration buffer (20 mM tris, 200 mM NaCl, 30 mM Imidazole; pH 8.0). Next, the column was washed with 20 volumes of equilibration buffer, followed by protein elution using a gradient of equilibration and elution buffer (20 mM Tris, 200 mM NaCl, 500 mM Imidazole; pH 8.0). Protein fractions were analysed on SDS-PAGE gels (Bio-Rad, California, USA) for purity and abundance. Suitable fractions were pooled and dialyzed against low salt solution (25 mM tris, 25 mM NaCl; pH 8.0) overnight at 4°C. Protein concentrations were determined using the Bradford method (Bradford, 1976).

### Bioassays on *Arabidopsis thaliana* and cotton seedlings

To assess the effect of treatment with a D effector protein and a *D-like1* protein from *Verticillium dahliae*, bioassays were conducted on *Arabidopsis thaliana* and cotton seedlings. For the *Arabidopsis thaliana* (Columbia-0) bioassay, ethanol-sterilized seeds were placed in 96-well plates with semi-solid media (½ MS, 0.2% agar, 0.5% sucrose) supplemented with the protein homologs to a final concentration of 0.2 mg/mL Bovine Serum Albumin (BSA) at the same concentration and the dialysis buffer alone (2.8 mM K₂HPO₄, 2.1 mM KH₂PO₄, 200 mM NaCl, pH 7.1) served as negative controls. The seedlings were stratified at 4°C in the dark for 24 hours and then incubated in a growth chamber at 22°C with a 16/8 hour day/night cycle for 14 days. For the cotton bioassay, one-week-old seedlings were uprooted, and the roots were placed into 10 mL protein solutions (5 µM). BSA (5 µM) was used as negative control, and the *V. dahliae* D effector protein (5 µM) was used as a positive control. The cotton seedlings were incubated in a growth chamber for 10 days at 22°C/19°C (day/night) with 70% relative humidity. In both experiments, plants were regularly monitored for phenotypic differences. At the end of the treatments, plants were photographed, and ImageJ was used to determine the canopy area (leaf area in cotton was measured at 2, 4, 7, and 10 days post-treatment). The data were analysed using an analysis of variance (ANOVA) combined with Tukey’s HSD (Honestly Significant Difference) test to identify significant differences.

### Calculation of Alien Indexes

Prior to calculating the alien index of genes in *Verticillium dahliae* strain CQ2, the gene models were improved from a previous annotation^38^, which was based solely on coding sequence prediction using ‘pipeline C’ of BRAKER version 3.0.8^139^ and failed to annotate the *D* gene. In this study, genes in the chromosomal-level assembly of CQ2^27^, in which repetitive elements were previously soft-masked^38^, were re-predicted using ‘pipeline D’ of the same BRAKER version with options ‘––gff3’ and ‘––fungus’, incorporating expression evidence from RNA-Seq data deposited in the NCBI Short Read Archive (accession numbers: SRR13442519, SRR13442520, and SRR13442521) that were generated in the previous study^140^. In addition, gene annotations from *V. dahliae* strain JR2 (accession number VDAG_JR2v.4.0 in EnsemblFungi)^33^ were transferred to CQ2 using GeMoMa version 1.9^141,142^, combined with a translated nucleotide-to-nucleotide search performed with MMseqs2 version 18.8cc5c^143^. All annotation datasets were then combined in a non-redundant manner using GffCompare version 0.12.10^113^, with the following priority order: BRAKER ‘pipeline D’ annotations, JR2-based ortholog annotations, and BRAKER ‘pipeline C’ annotations. The combined annotation was finalized using the same GeMoMa version to assign gene names according to their genomic positions. The TEs in *V. dahliae* CQ2 were annotated previously^38^, in which Helitron-like elements were identified with HelitronScanner^144^ in the TE annotation pipeline EDTA version 2.2.1^125^.

The alien index (AI) was calculated using the previously established formula AI = log((best E-value for ingroup) + e-200) – log((best E-value for outgroup) + e-200)^34^. E-values for ingroup and outgroup hits were obtained through nucleotide-to-nucleotide similarity search with Basic Local Alignment Search Tool (BLAST) version 2.16.0+, using the ‘blastn’ command with options ‘-word_size 11’, ‘-evalue 0.05’ and ‘-outfmt 6’ with default output plus query coverage (qcovs). Genes and TEs located from the start to 1,200,000 bp on chromosome 4 (Chr4) of the CQ2 genome (https://doi.org/10.5281/zenodo.15450312) assembly were used, and only hits covering more than 50% of the query sequences (qcovs > 50) and with an E-value lower than 0.05 were retained for alien index calculation. The E-value for a no hit was defined as 1 when either the ingroup or the outgroup had hits. AI values were not determined when both the ingroup and the outgroup had no hits. The coordinates of the queries are listed in Table S5 with AI values. The subject genome database was constructed using all currently available Pezizomycotina genome assemblies, downloaded along with taxonomy data from the NCBI database under the accession numbers listed in Table S6, with classification of ingroup and outgroup. Assemblies without order-level assignments were excluded. Downloads were performed using NCBI Datasets with the option ‘taxon 147538’. For the statistical analysis of the alien index (AI), genes located between 588,500 bp and 1,029,595 bp on Chr4 were designated as *Ar1h2 Starship* cargo, while the remaining genes in the core regions, excluding AGRs assigned in the previous study^38^, from the start of the chromosome to 1,200,000 bp were considered core genes. The normality and homoscedasticity of data were tested by the Shapiro–Wilk test and F test, respectively, with the R base package ‘stats’ version 4.4.2^106^, which indicated that the AI values were non-normally distributed and exhibited unequal variance (*p* < 0.01). The statistical difference in AI values between the two groups was assessed using the Brunner-Munzel test with the R package ‘lawstat’^145^.

### Circular dichroism spectroscopy

Circular dichroism spectroscopy was performed in the far-UV region using a Jasco J710 instrument (Jasco Inc., Easton, MD, USA) equipped with a Peltier apparatus at 20°C. Three spectra were collected using 0.2 mg/mL μM proteins in 20 mM Tris-HCl pH 7.5 and 150 mM NaCl buffer using a 1 mm quartz cell with a 50 nm/min scanning speed and averaged.

### Size exclusion chromatography

Size exclusion chromatography was performed on a Superdex75 10/300 column (GE Healthcare, Chicago, Illinois, USA). D*Fov* (1 mg/mL) was loaded on the column equilibrated with 20 mM Tris-HCl pH 7.5, 150 mM NaCl and connected to an HLPC AZURA system (KNAUER, Berlin, Germany). The injection volume was 500 μL, and the flow rate was 1 mL/min. The calibration curve was obtained by running the following protein standards: albumin (66.5 kDa), ovalbumin (43 kDa), chymotrypsinogen (25 kDa) and ribonuclease A (13.7 kDa). The protein eluted with a Ve = 11.7 ml corresponding to an apparent molecular weight of 35 kDa.

### Crystallization conditions and structure determination

Prior to crystallization, D*Fov* was further purified by size exclusion chromatography on a Superdex75 10/300 column (GE Healthcare, Chicago, Illinois, USA) equilibrated with 20 mM Tris-HCl pH 7.5, 150 mM NaCl. Crystallization conditions were initially screened automatically using the Oryx 4 crystallization platform (Douglas Instruments, East Garston, Berkshire, UK) by mixing equal amounts (0.4 µL) of reservoir solution and protein at two different concentrations (8.0 and 3.9 mg/mL in the SEC elution buffer). Small crystals were observed in the D2 condition of the Crystal Screen (Hampton Research, Aliso Viejo, CA, USA) after one week of incubation and further optimized by hanging drop vapour diffusion method. The best diffracting crystals were obtained by mixing 1 µL of protein solution (7.6 mg/mL) with 1 µl of reservoir (0.1 M Hepes pH 7.5, 1.4 M Sodium Citrate tribasic dihydrate) equilibrated versus 500 µL of the reservoir at 21°C. Crystals were transferred into a mother liquor solution containing 1 mM HgCl_2_ for 12 h, then transferred into another drop containing the mother liquid and 20% glycerol for 20 s before flash-freezing in liquid N_2_. X-ray diffraction data were collected at the ESRF Synchrotron in Grenoble (FR) on the beamline ID23-1. A complete data set was collected at a wavelength of 0.992 Å. Data were processed with XDS^146^, while analysis were performed with the autoPROC toolbox ^147^ and scaled with Aimless ^148^ at a final resolution of 1.47 Å in the CCP4 suite ^149^. Space group was determined based on systematic absences with Pointless^150^, and was P 1 2_1_ 1 with the following cell parameters (a, b, c Å – α, β, γ °): 27.8, 50.0, 72.3 – 90.0, 98.9, 90.0. The Matthews coefficient was 1.85 Å^3^Da^−1^, corresponding to one molecule per asymmetric unit and a solvent content of 33.5%. Experimental phasing was carried out with single-wavelength anomalous diffraction based on Hg anomalous scattering by the automatic software Crank-2^151^ as implemented in CCP4-cloud suite^149^. Iterative cycles of model building and refinement were performed with Coot ^152^ and Refmac5^153^. The final model consists of 77 protein residues (1493 non-H atoms), one Hepes (4-(2-hydroxyethyl)-1-piperazine ethanesulfonic acid) molecule, three Hg atoms and 184 water molecules. The full statistics for data collection and refinement are reported in Table S9. Coordinates and structure factors have been deposited in the Protein data bank with accession code 9FEI.

### Structural relationships and network rendering

The database of effector protein structures^52^ was downloaded from https://zenodo.org/records/7506581 and the D*Fov* protein structure was added after which structural similarities were calculated as previously described^52^.

## ACKNOWLEDGEMENTS

AD, JL and GLF acknowledge PhD fellowships from the University of Rome “La Sapienza”, China Scholarship Council (CSC) and the Coordination for the Improvement of Higher Education Personnel (CAPES) from the federal government of Brazil, respectively. Y.S. acknowledges funding through an Overseas Research Fellowship from the Japan Society for the Promotion of Science. TL and LFZ acknowledge funding by the National Natural Science Foundation of China (32372499 and 32230076, respectively). LF acknowledges financial support under the National Recovery and Resilience Plan (NRRP), Mission 4, Component 2, Investment 1.1, Call for tender No. 104 published on 2.2.2022 by the Italian Ministry of University and Research (MUR), funded by the European Union – NextGenerationEU– Project Title TAKLPAT – CUP E53D23007610006 – Grant Assignment Decree No. 104 adopted on 02-02-2022 by the Italian Ministry of Ministry of University and Research (MUR). BPHJT acknowledges funding by the Alexander von Humboldt Foundation in the framework of an Alexander von Humboldt Professorship endowed by the German Federal Ministry of Education and is furthermore supported by the Deutsche Forschungsgemeinschaft (DFG, German Research Foundation) under Germanýs Excellence Strategy – EXC 2048/1 – Project ID: 390686111 and the Research Council for Earth and Life Sciences (ALW) of the Netherlands Organization for Scientific Research (NWO). Bert Essenstam (Unifarm) is thanked for excellent plant care.

## AUTHOR CONTRIBUTIONS

LF and BPHJT conceived the project. AD, GLF, JL, MFS, LF, and BPHJT designed the experiments. AD, GLF, JL, IZ, YS, GL, WG, GG and GCMB performed the experiments and analysed the data. CGC, AV and JM performed the experiments on olive plants. BZ, LZ, TL and HT provided the *V. dahliae* strains and performed initial tests on cotton plants. MR provided the *Fusarium* strains used in this work. FT, MCBDP, ADM, GG solved the crystal structure of D*Fov*. AD, GLF, JL, YS, LF and BPHJT wrote the manuscript. All authors read and approved the final manuscript.

## DATA AVAILABILITY

*D-like* sequences are deposited at Zenodo (https://doi.org/10.5281/zenodo.18375323). The chromosomal-level assemblies of *Verticillium* strains CQ2, VLB2, and PD683 are available on Zenodo (https://doi.org/10.5281/zenodo.15450312). The other genome assemblies used in this study are available on NCBI under the accession numbers listed in the Supplementary Tables.

## SUPPORTING INFORMATION

**Figure S1.** Phylogenetic tree of in house-sequenced *V. dahliae* strains. *V. dahliae* strains clustering with known D pathotype strains (TM6, V991, V117 and T9) are showed in lightred, while strains clustering with known ND pathotype strains (V4, BP2, cd3 and HN) are showed in light-green. Strains that distinct from the D and ND pathotype groups are showed in light-blue. Strains that used for phenotypic characterization in this work are display in bold with blue color. Phylogenetic relationship between sequenced *V. dahliae* strains was inferred using RealPhy (Bertels et al., 2014) and *V. dahliae* strain JR2 used as reference. *V. alfalfae* ms102 was used as root of the tree.

**Figure S2.** Clustering analysis of seventy-two sequenced *V. dahliae* strains. Sequences of seventy-two *V. dahliae* strains were aligned on the assembled genome of CQ2 and clustered in three groups based on presence/absence polymorphism. Strains clustering with known D pathotype strains are displayed in blue, while strains clustering with ND pathotype strains are showed in red. Strain BGI_32 that is the most divergent from the two groups showed in black.

**Figure S3.** Phenotype of cotton plants inoculated with *V. dahliae* strain BGI32. (A) Typical phenotype of cotton plants (cv. Xinluzao63) upon mock-inoculation or inoculation with BP2, BGI32 and CQ2 at 28 days post inoculation (dpi). ND pathotype strain BP2 and D pathotype strain CQ2 were used as inoculation controls. (B) Defoliation was classified as 0 (0% leaf drop off), 1 (<25% leaf drop off), 2 (<50% leaf drop off), 3 (<75% leaf drop off) and 4 (<100% leaf drop off). Inoculation experiments were performed with ten plants for each fungal strain and repeated twice independently with similar results.

**Figure S4.** Expression analysis of the seven candidate genes. The heatmap showed the expression level of each gene during a time course of cotton infected by D pathotype strain V991 at 6, 9, 12 and 15 days post inoculation (DPI). The scaled expression values are color-coded according to the scale bar in the left top corner.

**Figure S5.** Construction and verification of *D* single and double deletion mutants. (A) Schematic representation of the homologous recombination events to establish targeted replacement of *D* gene with phosphotransferase (HPH) and the nourseothricin resistance gene cassette (NAT). (B) Verification of D single deletion (Δ*D*) and double deletion mutants (ΔΔ*D*) in *V. dahliae* cotton defoliating strain CQ2 by PCR. Amplicons generated with outside primers indicated in panel A are shown for wild type strain CQ2 (WT), two Δ*D* mutants (#1 and #2) and two ΔΔ*D* mutants (#1 and #2). (C) Verification of *D* single deletion (Δ*D*) and double deletion mutants (ΔΔ*D*) in *V. dahliae* olive defoliating strain V150I by PCR. Amplicons generated with outside primers indicated in panel A are shown for wild-type strain V150I (WT), one Δ*D* mutant (#7) and three ΔΔ*D* mutants (#5, #7 and #8).

**Figure S6.** Detection of *D* gene transcripts in various *D* deletion and complementation strains. Amplification of *D* gene fragment (from left to right) from cDNA in wild type strain CQ2 (WT), two Δ*D* strains (#1 and #2), two Δ*D* complementation strains (Δ*D*/p*D::D* #1 and #2), two ΔΔ*D* strains (#1 and #2) and one ΔΔ*D* complementation strain (ΔΔ*D*/p*D*::*D* #3). *V. dahliae GAPDH* gene was used as endogenous control.

**Figure S7.** Expression of *D* gene (p*D*::*D*) in ND pathotype strain JR2. Amplification of *D* gene fragment (from left to right) from cDNA in wild type strain JR2, one *D* expression (p*D::D*) transformant of JR2 and CQ2 (used as positive control). *V. dahliae GAPDH* gene was used as endogenous control.

**Figure S8.** Introduction of *D* gene in ND pathotype strain results in defoliation symptoms. (A) Typical phenotype of cotton (cv. Xinluzao63) plants that were mock-inoculated or inoculated with ND pathotype strain JR2, one *D* expression transformant of JR2 (JR2 p*D*::*D*) and D pathotype strain CQ2 at 28 days post inoculation (dpi). (B) Defoliation was classified as 0 (0% leaves drop off), 1 (<25% leaves drop off), 2 (<50% leaves drop off), 3 (<75% leaves drop off) and 4 (<100% leaves drop off) at 28 dpi. (C) Fungal biomass as determined with real-time PCR at 28 dpi. Bars indicate the *V. dahliae* biomass relatively to the cotton biomass. Significant differences were calculated with the Mann-Whitney U test (P < 0.05) and depicted by different letter labels. Error bars represent the standard error.

**Figure S9.** D effector protein induces cotton defoliation *in vitro*. (A-D) Typical appearance of cotton seedlings (cv. Xinluzao63) treated with D effector protein at 10 days (A-B) and 16 days (C-D). Note, marginal chlorosis and wilting symptoms on cotyledons at 10 days (B) and severe chlorosis, and leaf drop off (red arrow; D). (E) Bars represent the average defoliation of two biological replicates with standard deviation. Experiments were repeated twice independently with similar results.

**Figure S10.** PCR detection of *D* gene in *V. dahliae* strains. Amplification of *D* gene fragment (from left to right) from genomic DNA in CQ2, V150I, V641I, V403I, V356I, 812I. As an endogenous control, a fragment of the *Verticillium* ITS region was amplified.

**Figure S11.** The D effector is responsible for olive defoliation. Typical phenotype of olive (cv. Picual) upon mock-inoculation or inoculation with wild type strain CV150I (WT), and three ΔΔ*D* mutants (ΔΔ*D* #5, ΔΔ*D* #7 and ΔΔ*D* #8) at 132 days post inoculation (dpi).

**Figure S12.** The *D-like 4* allele is disrupted by a retrotransposon of the family Ty3/Gypsy-like. Schematic representation of the *D-like4* allele, showing the interruption of its sequence (represented in green) by a retrotransposon of the family Ty3/Gypsy-like (represented in grey). Only a short portion (174 bp) of the retrotransposon is present before the end of the contig in which the D-like 4 allele is present in both *V. dahliae* strains that carry this allele (V574 and V700).

**Fig. S13.** *D* gene horizontal transfer. (A) Maximum-likelihood phylogenetic tree of *D* homologs inferred using a nucleotide substitution model. Scale bars indicate nucleotide substitutions per site. Circles at nodes denote bootstrap support values >95% based on 1,000 replicates. (B) Rates of synonymous substitutions per synonymous site (*K*_s_) between species. The *D* homologs selected for testing are indicated by red arrows in (A). Glyceraldehyde 3-phosphate dehydrogenase (*GAPDH*) genes from the same strains were used as vertically inherited core housekeeping gene references.

**Fig. S14.** Distribution of homologs of *Ar1h2 Starship* cargo genes and transposable elements (TEs) across Pezizomycotina. Cargo elements from the *Ar1h2 Starship* in *V. dahliae* strain CQ2 were used as queries. Circles indicate the coverage and sequence identity of the best hits in each genus within the orders Glomerellales and Hypocreales, as well as in all other genera. Best hits across all genera excluding *Verticillium* are outlined in red.

**Figure S15.** Circular dichroism spectroscopy and size exclusion chromatography analysis of the D*Fov* protein. (A) Far-UV CD spectra of *E. coli*-produced D*Fov* protein (0.2 mg/ml) in 20 mM Tris-HCl pH 7.5, 150 mM NaCl buffer. The spectrum reveals that the recombinant protein is well folded with a prevalently beta-sheet secondary structure. (B) Size exclusion chromatographic elution profile of purified D*Fov*. The protein, loaded on a Superdex 75 10/300 column, eluted as a single sharp peak at 11.7 ml, corresponding to an apparent molecular weight of 35 kDa. Considering that the purified protein, including the affinity tag, has a predicted Mw of 26.6 kDa, this value indicates that D*Fov* is monomeric and displays an elongated shape. *Inset*: the column was calibrated by running the following protein standards: albumin (66.5 kDa), ovalbumin (43 kDa), chymotrypsinogen (25 kDa) and ribonuclease A (13.7 kDa).

**Figure S16.** The AlphaFold3-predicted D*Fov* model diverges from the D*Fov* protein structure determined with X-ray crystallography. (A) D*Fov* crystal structure, determined by X-ray crystallography. (B) D*Fov* model predicted with AlphaFold3. (C) Structural alignment of the D*Fov* crystal structure and the AlphaFold3 model. TM-score indicates topological similarity between two protein structures reported on a scale from 0 to 1, where 1 indicates a perfect match, while the RMSD score indicates the average deviation between the corresponding atoms of two proteins with values lower than 2 indicating significant similarity between structures.

**Figure S17.** D*Fov* is a structurally unique effector. (A) Structural similarity network based on pairwise DALI Z-scores. Each dot represents a protein structure while different colors represent different structural families. The D*Fov* protein structure is indicated with as well as effector proteins that are somewhat similar, including Alt-A1, Tox3, Zuotin and FSA2. (B) D*Fov*, Tox3 and Alt-A1 protein structures.

## REFENCES

1. Zhang, X. et al. A panoramic view of cotton resistance to Verticillium dahliae: From genetic architectures to precision genomic selection. iMeta 4, (2025).

2. Fradin, E. F. & Thomma, B. P. H. J. Physiology and molecular aspects of Verticillium wilt diseases caused by *V. dahliae* and *V. albo-atrum*. Mol. Plant Pathol. 7, 71–86 (2006).

3. Klosterman, S. J., Atallah, Z. K., Vallad, G. E. & Subbarao, K. V. Diversity, pathogenicity, and management of verticillium species. Annu. Rev. Phytopathol. 47, 39–62 (2009).

4. Inderbitzin, P. & Subbarao, K. V. Verticillium systematics and evolution: How confusion impedes verticillium wilt management and how to resolve it. Phytopathology 104, 564–574 (2014).

5. Schnathorst, W. & Mathre, D. Host range and differentiation of a severe form of *Verticillium albo-atrum* in cotton. Phytopathology 56, 1155–1161 (1966).

6. Schnathorst WC & Sibbet GS. TI VERTICILLIUM STRAIN… a Major Factor in Cotton and Olive Wilt. (1971).

7. Jiménez-Díaz, R. M., Olivares-García, C., Landa, B. B., Del Mar Jiménez-Gasco, M. & Navas-Cortés, J. A. Region-wide analysis of genetic diversity in *Verticillium dahliae* populations infecting olive in southern Spain and agricultural factors influencing the distribution and prevalence of vegetative compatibility groups and pathotypes. Phytopathology 101, 304–315 (2011).

8. Korolev, N., et al. Vegetative compatibility of cotton-defoliating *Verticillium dahliae* in Israel and its pathogenicity to various crop plants. Eur. J. Plant Pathol. 122, 603–617 (2008).

9. Dervis, S., Mercado-Blanco, J., Erten, L., Valverde-Corredor, A. & Pérez-Artés, E. Verticillium wilt of olive in Turkey: A survey on disease importance, pathogen diversity and susceptibility of relevant olive cultivars. Eur. J. Plant Pathol. 127, 287–301 (2010).

10. López-Escudero, F. J., Del Río, C., Caballero, J. M. & Blanco-López, M. A. Evaluation of olive cultivars for resistance to *Verticillium dahliae*. Eur. J. Plant Pathol. 110, 79–85 (2004).

11. Milgroom, M. G., Jiménez-Gasco, M. D. M., Olivares-García, C. & Jiménez-Díaz, R. M. Clonal expansion and migration of a highly virulent, defoliating lineage of *Verticillium dahliae*. Phytopathology 106, 1038–1046 (2016).

12. Jiménez-Díaz, R. M., et al. Variation of pathotypes and races and their correlations with clonal lineages in *Verticillium dahliae*. Plant Pathol. 66, 651–666 (2017).

13. Pérez-Artés, E., García-Pedrajas, M. D., Bejarano-Alcázar, J. & Jiménez-Díaz, R. M. Differentiation of cotton-defoliating and nondefoliating pathotypes of *Verticillium dahliae* by RAPD and specific PCR analyses. Eur. J. Plant Pathol. 106, 507–517 (2000).

14. Mercado-Blanco, J., Rodríguez-Jurado, D., Pérez-Artés, E. & Jiménez-Díaz, R. M. Detection of the defoliating pathotype of *Verticillium dahliae* in infected olive plants by nested PCR. Eur. J. Plant Pathol. 108, 1–13 (2002).

15. Collins, A., et al. Correlation of molecular markers and biological properties in *Verticillium dahliae* and the possible origins of some isolates. Plant Pathol. 54, 549–557 (2005).

16. Rovenich, H., Boshoven, J. C. & Thomma, B. P. H. J. Filamentous pathogen effector functions: Of pathogens, hosts and microbiomes. Curr. Opin. Plant Biol. 20, 96–103 (2014).

17. Cook, D. E., Mesarich, C. H. & Thomma, B. P. H. J. Understanding Plant Immunity as a Surveillance System to Detect Invasion. Annual Review of Phytopathology vol. 53 (2015).

18. Snelders, N. C., et al. Microbiome manipulation by a soil-borne fungal plant pathogen using effector proteins. Nat. Plants 6, 1365–1374 (2020).

19. Snelders, N. C., Petti, G. C., van den Berg, G. C. M., Seidl, M. F. & Thomma, B. P. H. J. An ancient antimicrobial protein co-opted by a fungal plant pathogen for in planta mycobiome manipulation. Proc. Natl. Acad. Sci. U. S. A. 118, (2021).

20. Snelders, N. C., et al. A highly polymorphic effector protein promotes fungal virulence through suppression of plant-associated Actinobacteria. New Phytologist 237, 944–958 (2023).

21. Kraege, A., et al. Undermining the cry for help: the phytopathogenic fungus *Verticillium dahliae* secretes an antimicrobial effector protein to undermine host recruitment of antagonistic *Pseudomonas* bacteria. New Phytologist 249, 406–417 (2026).

22. Mesny, F., Bauer, M., Zhu, J. & Thomma, B. P. H. J. Meddling with the microbiota: Fungal tricks to infect plant hosts. Curr. Opin. Plant Biol. 82, 102622 (2024).

23. Gibriel, H. A. Y., Thomma, B. P. H. J. & Seidl, M. F. The age of effectors: Genome-based discovery and applications. Phytopathology 106, 1206–1212 (2016).

24. Xu, F., et al. Prevalence of the defoliating pathotype of *Verticillium dahliae* on cotton in central China and virulence on selected cotton cultivars. Journal of Phytopathology 160, 369–376 (2012).

25. Zhang, B., et al. Island cotton Gbve1 gene encoding a receptor-like protein confers resistance to both defoliating and non-defoliating isolates of *Verticillium dahliae*. PLoS One 7, e51091 (2012).

26. Liu, T., et al. Unconventionally secreted effectors of two filamentous pathogens target plant salicylate biosynthesis. Nat. Commun. 5, 4686 (2014).

27. Seidl, M. F., et al. Repetitive elements contribute to the diversity and evolution of centromeres in the fungal genus Verticillium. mBio 11, 1–22 (2020).

28. Depotter, J. R. L., et al. Dynamic virulence-related regions of the plant pathogenic fungus *Verticillium dahliae* display enhanced sequence conservation. Mol. Ecol. 28, 3482–3495 (2019).

29. Finn, R. D. et al. InterPro in 2017—beyond protein family and domain annotations. Nucleic Acids Res. 45, D190–D199 (2017).

30. Kombrink, A., et al. *Verticillium dahliae* LysM effectors differentially contribute to virulence on plant hosts. Mol. Plant Pathol. 18, 596–608 (2017).

31. Maldonado-González, M. M., Bakker, P. A. H. M., Prieto, P. & Mercado-Blanco, J. Arabidopsis thaliana as a tool to identify traits involved in *Verticillium dahliae* biocontrol by the olive root endophyte *Pseudomonas fluorescens* PICF7. Front. Microbiol. 6, (2015).

32. Collado-Romero, M., Mercado-Blanco, J., Olivares-García, C., Valverde-Corredor, A. & Jiménez-Díaz, R. M. Molecular variability within and among *Verticillium dahliae* vegetative compatibility groups determined by fluorescent amplified fragment length polymorphism and polymerase chain reaction markers. Phytopathology 96, 485–495 (2006).

33. Faino, L., et al. Single-molecule real-time sequencing combined with optical mapping yields completely finished fungal genome. mBio 6, (2015).

34. Gladyshev, E. A., Meselson, M. & Arkhipova, I. R. Massive horizontal gene transfer in bdelloid rotifers. Science (1979). 320, 1210–1213 (2008).

35. Bucknell, A. H. & McDonald, M. C. That’s no moon, it’s a *Starship*: Giant transposons driving fungal horizontal gene transfer. Mol. Microbiol. 120, 555–563 (2023).

36. Urquhart, A., Vogan, A. A. & Gluck-Thaler, E. Starships: a new frontier for fungal biology. Trends in Genetics 40, 1060–1073 (2024).

37. Urquhart, A. S., Gluck-Thaler, E. & Vogan, A. A. Gene acquisition by giant transposons primes eukaryotes for rapid evolution via horizontal gene transfer. Sci. Adv. 10, (2024).

38. Sato, Y., et al. Starship giant transposons dominate plastic genomic regions in a fungal plant pathogen and drive virulence evolution. Nat. Commun. 16, 6806 (2025).

39. Urquhart, A. S., Forsythe, A. & Vogan, A. A. Are fungal disease outbreaks instigated by *Starship* transposons? Mol. Plant Pathol. 26, (2025).

40. Gluck-Thaler, E. & Vogan, A. A. Systematic identification of cargo-mobilizing genetic elements reveals new dimensions of eukaryotic diversity. Nucleic Acids Res. 52, 5496–5513 (2024).

41. Jobe, T. O., Ulloa, M. & Ellis, M. L. A high-quality whole-genome sequence, assembly, and gene annotation of *Fusarium oxysporum* f. sp. *vasinfectum* (Fov) race 1 from California. Microbiol. Resour. Announc. 13, (2024).

42. Komissarov, E. N., et al. Genomic differences between two *Fusarium oxysporum formae speciales* causing root rot in cucumber. Journal of Fungi 11, 140 (2025).

43. Jobe, T. O., Ulloa, M. & Ellis, M. L. Two *de novo* genome assemblies from pathogenic *Fusarium oxysporum* f. sp. *vasinfectum* race 4 (FOV4) isolates from California. Microbiol. Resour. Announc. 13, (2024).

44. van Dam, P., et al. A mobile pathogenicity chromosome in *Fusarium oxysporum* for infection of multiple cucurbit species. Sci. Rep. 7, 9042 (2017).

45. Gluck-Thaler, E., et al. Giant *Starship* Elements Mobilize Accessory Genes in Fungal Genomes. Mol. Biol. Evol. 39, (2022).

46. Kapitonov, V. V. & Jurka, J. Rolling-circle transposons in eukaryotes. Proceedings of the National Academy of Sciences 98, 8714–8719 (2001).

47. Grabundzija, I., et al. A Helitron transposon reconstructed from bats reveals a novel mechanism of genome shuffling in eukaryotes. Nat. Commun. 7, 10716 (2016).

48. Barro-Trastoy, D. & Köhler, C. Helitrons: genomic parasites that generate developmental novelties. Trends in Genetics 40, 437–448 (2024).

49. Abramson, J., et al. Accurate structure prediction of biomolecular interactions with AlphaFold 3. Nature 630, 493–500 (2024).

50. Holm, L. Dali server: structural unification of protein families. Nucleic Acids Res. 50, W210–W215 (2022).

51. Outram, M. A., et al. The crystal structure of SnTox3 from the necrotrophic fungus *Parastagonospora nodorum* reveals a unique effector fold and provides insight into Snn3 recognition and pro-domain protease processing of fungal effectors. New Phytologist 231, 2282–2296 (2021).

52. Derbyshire, M. C. & Raffaele, S. Surface frustration re-patterning underlies the structural landscape and evolvability of fungal orphan candidate effectors. Nat. Commun. 14, 5244 (2023).

53. Jones, J. D. G. & Dangl, J. L. The plant immune system. Nature 444, 323–329 (2006).

54. De Jonge, R., Bolton, M. D. & Thomma, B. P. H. J. How filamentous pathogens co-opt plants: The ins and outs of fungal effectors. Curr. Opin. Plant Biol. 14, 400–406 (2011).

55. Jones, J. D. G., Staskawicz, B. J. & Dangl, J. L. The plant immune system: From discovery to deployment. Cell 187, 2095–2116 (2024).

56. Swarup, S. A Pathogenicity Locus from *Xanthomonas citri* Enables Strains from Several Pathovars of *X. campestris* to Elicit Cankerlike Lesions on Citrus. Phytopathology 81, 802 (1991).

57. Swarup, S. An *Xanthomonas citri* pathogenicity gene, *pthA,* pleiotropically encodes gratuitous avirulence on nonhosts. Molecular Plant-Microbe Interactions 5, 204 (1992).

58. Duan, Y. P., Castañeda, A., Zhao, G., Erdos, G. & Gabriel, D. W. Expression of a Single, Host-Specific, Bacterial Pathogenicity Gene in Plant Cells Elicits Division, Enlargement, and Cell Death. Molecular Plant-Microbe Interactions® 12, 556–560 (1999).

59. Liu, Z., et al. SnTox1, a *Parastagonospora nodorum* necrotrophic effector, is a dual-function protein that facilitates infection while protecting from wheat-produced chitinases. New Phytologist 211, 1052–1064 (2016).

60. Bhat, R. G. & Subbarao, K. V. Host range specificity in *Verticillium dahliae*. Phytopathology 89, 1218–1225 (1999).

61. Chen, J. Y., et al. Comparative genomics reveals cotton-specific virulence factors in flexible genomic regions in *Verticillium dahliae* and evidence of horizontal gene transfer from Fusarium. New Phytologist 217, 756–770 (2018).

62. Usami, T., Itoh, M. & Amemiya, Y. Mating type gene MAT1-2-1 is common among Japanese isolates of *Verticillium dahliae*. Physiol. Mol. Plant Pathol. 73, 133–137 (2008).

63. Usami, T., Itoh, M. & Amemiya, Y. Asexual fungus *Verticillium dahliae* is potentially heterothallic. Journal of General Plant Pathology 75, 422–427 (2009).

64. Atallah, Z. K., et al. Population analyses of the vascular plant pathogen *Verticillium dahliae* detect recombination and transcontinental gene flow. Fungal Genetics and Biology 47, 416–422 (2010).

65. Milgroom, M. G., Jiménez-Gasco, M. del M., Olivares García, C., Drott, M. T. & Jiménez-Díaz, R. M. Recombination between clonal lineages of the asexual fungus Verticillium dahliae detected by genotyping by sequencing. PLoS One 9, e106740 (2014).

66. Rafiei, V., et al. Comparison of genotyping by sequencing and microsatellite markers for unravelling population structure in the clonal fungus *Verticillium dahliae*. Plant Pathol. 67, 76–86 (2018).

67. Joaquim, T. R. & Rowe, R. C. Reassessment of vegetative compatibility relationships among strains of Verticillium dahliae using nitrate-nonutilizing mutants. Reactions 4, 10–1094 (1990).

68. Strausbaugh, C. A. Assessment of vegetative compatibility and virulence. Phytopathology 83, 1253–1258 (1993).

69. Daayf, F., Nicole, M. & Geiger, J.-P. Differentiation of *Verticillium dahliae* populations on the basis of vegetative compatibility and pathogenicity on cotton. Eur. J. Plant Pathol. 101, 69–79 (1995).

70. Hiemstra, J. A. & Rataj-Guranowska, M. Vegetative compatibility groups in *Verticillium dahliae* isolates from the Netherlands as compared to VCG diversity in Europe and in the USA. Eur. J. Plant Pathol. 109, 827–839 (2003).

71. Bell, A. A. Verticillium wilt. in Cotton Diseases (ed. Hillocks, R. J.) 87–126 (CAB International, Wallingford, 1992).

72. Friesen, T. L., et al. Emergence of a new disease as a result of interspecific virulence gene transfer. Nat. Genet. 38, 953–956 (2006).

73. Raffaele, S., et al. Genome evolution following host jumps in the irish potato famine pathogen lineage. Science (1979). 330, 1540–1543 (2010).

74. Stukenbrock, E. H., Christiansen, F. B., Hansen, T. T., Dutheil, J. Y. & Schierup, M. H. Fusion of two divergent fungal individuals led to the recent emergence of a unique widespread pathogen species. Proceedings of the National Academy of Sciences 109, 10954–10959 (2012).

75. Dutheil, J. Y., et al. A tale of genome compartmentalization: the evolution of virulence clusters in smut fungi. Genome Biol. Evol. 8, 681–704 (2016).

76. De Jonge, R., et al. Tomato immune receptor Ve1 recognizes effector of multiple fungal pathogens uncovered by genome and RNA sequencing. Proc. Natl. Acad. Sci. U. S. A. 109, 5110–5115 (2012).

77. De Jonge, R., et al. Extensive chromosomal reshuffling drives evolution of virulence in an asexual pathogen. Genome Res. 23, 1271–1282 (2013).

78. Faino, L., et al. Transposons passively and actively contribute to evolution of the two-speed genome of a fungal pathogen. Genome Res. 26, 1091–1100 (2016).

79. Shen, D., et al. Gene duplication and fragment recombination drive functional diversification of a superfamily of cytoplasmic effectors in *Phytophthora sojae*. PLoS One 8, e70036 (2013).

80. Pennington, H. G., et al. The fungal ribonuclease-like effector protein CSEP0064/BEC1054 represses plant immunity and interferes with degradation of host ribosomal RNA. PLoS Pathog. 15, e1007620 (2019).

81. Cao, Y., et al. Structural polymorphisms within a common powdery mildew effector scaffold as a driver of coevolution with cereal immune receptors. Proceedings of the National Academy of Sciences 120, (2023).

82. Chavarro-Carrero, E. A., et al. Comparative genomics reveals the *in planta-* secreted *Verticillium dahliae* Av2 effector protein recognized in tomato plants that carry the *V2* resistance locus. Environ. Microbiol. 23, 1941–1958 (2021).

83. Ma, L. J., et al. Comparative genomics reveals mobile pathogenicity chromosomes in Fusarium. Nature 464, 367–373 (2010).

84. Habig, M., Patneedi, S. K., Stam, R. & De Fine Licht, H. H. Horizontal transfer of accessory chromosomes in fungi – a regulated process for exchange of genetic material? Heredity (Edinb). https://doi.org/10.1038/s41437-025-00746-0 (2025) doi:10.1038/s41437-025-00746-0.

85. Sandell, L., et al. The role of toxin/antidote genes in the maintenance and evolution of accessory chromosomes in *Fusarium*. Genetics 231, (2025).

86. van Westerhoven, A. C., et al. Frequent genetic exchanges revealed by a pan-mitogenome graph of a fungal plant pathogen. mBio 15, (2024).

87. van Westerhoven, A. C., et al. Reference-free identification and pangenome analysis of accessory chromosomes in a major fungal plant pathogen. NAR Genom. Bioinform. 7, (2025).

88. McDonald, M. C., et al. Transposon-mediated horizontal transfer of the host-specific virulence protein ToxA between three fungal wheat pathogens. mBio 10, (2019).

89. Bucknell, A., et al. *Sanctuary*: a *Starship* transposon facilitating the movement of the virulence factor ToxA in fungal wheat pathogens. mBio 16, (2025).

90. Gourlie, R., McDonald, M. C., Hafez, M. & Aboukhaddour, R. The virulence gene ToxB is both amplified and disrupted by transposons in the wheat pathogen Pyrenophora tritici-repentis. Preprint at 10.1101/2025.11.13.688278 (2025).

91. Subieta, K. et al. A single pathogen-secreted protein reprograms plants for drought resilience. Preprint at 10.1101/2025.01.09.632073 (2025).

92. López-Escudero, F. J. & Mercado-Blanco, J. Verticillium wilt of olive: a case study to implement an integrated strategy to control a soil-borne pathogen. Plant Soil 344, 1–50 (2011).

93. Shaban, M., et al. Physiological and molecular mechanism of defense in cotton against *Verticillium dahliae*. Plant Physiology and Biochemistry 125, 193–204 (2018).

94. Laugé, R., et al. Successful search for a resistance gene in tomato targeted against a virulence factor of a fungal pathogen. Proceedings of the National Academy of Sciences 95, 9014–9018 (1998).

95. Takken, F. L. W., et al. A second gene at the tomato Cf-4 locus confers resistance to *Cladosporium fulvum* through recognition of a novel avirulence determinant. The Plant Journal 20, 279–288 (1999).

96. Vleeshouwers, V. G. A. A. & Oliver, R. P. Effectors as tools in disease resistance breeding against biotrophic, hemibiotrophic, and necrotrophic plant pathogens. Molecular Plant-Microbe Interactions 27, 196–206 (2014).

97. Vleeshouwers, V. G. A. A., et al. Understanding and exploiting late blight resistance in the age of effectors. Annu. Rev. Phytopathol. 49, 507–531 (2011).

98. Bertels, F., Silander, O. K., Pachkov, M., Rainey, P. B. & Van Nimwegen, E. Automated reconstruction of whole-genome phylogenies from short-sequence reads. Mol. Biol. Evol. 31, 1077–1088 (2014).

99. Langmead, B. & Salzberg, S. L. Fast gapped-read alignment with Bowtie 2. Nat. Methods 9, 357–359 (2012).

100. Stamatakis, A. RAxML version 8: a tool for phylogenetic analysis and post-analysis of large phylogenies. Bioinformatics 30, 1312–1313 (2014).

101. Gao, W., et al. Proteomic and Virus-induced Gene Silencing (VIGS) analyses reveal that gossypol, brassinosteroids, and jasmonic acid contribute to the resistance of cotton to verticillium dahliae. Molecular and Cellular Proteomics 12, 3690–3703 (2013).

102. Chin, C. S., et al. Nonhybrid, finished microbial genome assemblies from long-read SMRT sequencing data. Nat. Methods 10, 563–569 (2013).

103. Holt, C. & Yandell, M. MAKER2: An annotation pipeline and genome-database management tool for second-generation genome projects. BMC Bioinformatics 12, (2011).

104. Li, H. & Durbin, R. Fast and accurate long-read alignment with Burrows-Wheeler transform. Bioinformatics 26, 589–595 (2010).

105. Quinlan, A. R. & Hall, I. M. BEDTools: a flexible suite of utilities for comparing genomic features. Bioinformatics 26, 841–842 (2010).

106. R Core Team. R: A Language and Environment for Statistical Computing. Preprint at https://www.R-project.org/ (2021).

107. Trapnell, C., et al. Transcript assembly and quantification by RNA-Seq reveals unannotated transcripts and isoform switching during cell differentiation. Nat. Biotechnol. 28, 511–515 (2010).

108. Camacho, C., et al. BLAST+: architecture and applications. BMC Bioinformatics 2009 10:1 10, 421– (2009).

109. Löytynoja, A. & Goldman, N. webPRANK: a phylogeny-aware multiple sequence aligner with interactive alignment browser. BMC Bioinformatics 2010 11:1 11, 579– (2010).

110. Robert, X. & Gouet, P. Deciphering key features in protein structures with the new ENDscript server. Nucleic Acids Res. 42, W320–W324 (2014).

111. Sánchez-Vallet, A., et al. Fungal effector Ecp6 outcompetes host immune receptor for chitin binding through intrachain LysM dimerization. Elife 2, (2013).

112. Slater, G. S. C. & Birney, E. Automated generation of heuristics for biological sequence comparison. BMC Bioinformatics 6, 31 (2005).

113. Pertea, G. & Pertea, M. GFF Utilities: GffRead and GffCompare. F1000Res. 9, 304 (2020).

114. Shen, W., Sipos, B. & Zhao, L. SeqKit2: A Swiss army knife for sequence and alignment processing. iMeta 3, (2024).

115. O’Leary, N. A., et al. Exploring and retrieving sequence and metadata for species across the tree of life with NCBI Datasets. Sci. Data 11, 732 (2024).

116. Katoh, K. & Standley, D. M. MAFFT Multiple Sequence Alignment Software Version 7: Improvements in Performance and Usability. Mol. Biol. Evol. 30, 772–780 (2013).

117. Capella-Gutiérrez, S., Silla-Martínez, J. M. & Gabaldón, T. trimAl: a tool for automated alignment trimming in large-scale phylogenetic analyses. Bioinformatics 25, 1972–1973 (2009).

118. Wong, T. et al. IQ-TREE 3: Phylogenomic Inference Software using Complex Evolutionary Models. Preprint at 10.32942/X2P62N (2025).

119. Kalyaanamoorthy, S., Minh, B. Q., Wong, T. K. F., von Haeseler, A. & Jermiin, L. S. ModelFinder: fast model selection for accurate phylogenetic estimates. Nat. Methods 14, 587–589 (2017).

120. Hoang, D. T., Chernomor, O., von Haeseler, A., Minh, B. Q. & Vinh, L. S. UFBoot2: Improving the Ultrafast Bootstrap Approximation. Mol. Biol. Evol. 35, 518–522 (2018).

121. Yu, G., Smith, D. K., Zhu, H., Guan, Y. & Lam, T. T. **ggtree**: an **R** package for visualization and annotation of phylogenetic trees with their covariates and other associated data. Methods Ecol. Evol. 8, 28–36 (2017).

122. Paradis, E. & Schliep, K. ape 5.0: an environment for modern phylogenetics and evolutionary analyses in R. Bioinformatics 35, 526–528 (2019).

123. Wang, D., Zhang, Y., Zhang, Z., Zhu, J. & Yu, J. KaKs_Calculator 2.0: A Toolkit Incorporating Gamma-Series Methods and Sliding Window Strategies. Genomics Proteomics Bioinformatics 8, 77–80 (2010).

124. van Dam, P. & Rep, M. The distribution of miniature impala elements and SIX genes in the Fusarium genus is suggestive of horizontal gene transfer. J. Mol. Evol. 85, 14–25 (2017).

125. Ou, S., et al. Benchmarking transposable element annotation methods for creation of a streamlined, comprehensive pipeline. Genome Biol. 20, 275 (2019).

126. Marçais, G., et al. MUMmer4: A fast and versatile genome alignment system. PLoS Comput. Biol. 14, e1005944 (2018).

127. Gu, Z., Gu, L., Eils, R., Schlesner, M. & Brors, B. *circlize* implements and enhances circular visualization in R. Bioinformatics 30, 2811–2812 (2014).

128. Hackl, T., Ankenbrand, M. J. & van Adrichem, B. gggenomes: A Grammar of Graphics for Comparative Genomics. CRAN: Contributed Packages Preprint at 10.32614/CRAN.package.gggenomes (2024).

129. Levy Karin, E., Mirdita, M. & Söding, J. MetaEuk—sensitive, high-throughput gene discovery, and annotation for large-scale eukaryotic metagenomics. Microbiome 8, 48 (2020).

130. Fradin, E. F., et al. Genetic dissection of Verticillium wilt resistance mediated by tomato Ve1. Plant Physiol. 150, 320–332 (2009).

131. Song, Y., et al. Transfer of tomato immune receptor Ve1 confers Ave1-dependent Verticillium resistance in tobacco and cotton. Plant Biotechnol. J. 16, 638–648 (2018).

132. Leyva-Pérez, M. de la O., et al. Tolerance of olive (*Olea europaea*) cv Frantoio to *Verticillium dahliae* relies on both basal and pathogen-induced differential transcriptomic responses. New Phytol. 217, 671–686 (2018).

133. Gómez Lama Cabanás, C., et al. Indigenous Pseudomonas spp. Strains from the Olive (Olea europaea L.) rhizosphere as effective biocontrol agents against Verticillium dahliae: From the host roots to the bacterial genomes. Front. Microbiol. 9, 297478 (2018).

134. Zhang, L., et al. The Verticillium-specific protein VdSCP7 localizes to the plant nucleus and modulates immunity to fungal infections. New Phytol. 215, 368–381 (2017).

135. Campbell, C. L. & Madden, L. V. Introduction to Plant Disease Epidemiology. John Wiley & Sons, New York. – References – Scientific Research Publishing. https://www.scirp.org/reference/referencespapers?referenceid=1799831 (1990).

136. Frandsen, R. J., Andersson, J. A., Kristensen, M. B. & Giese, H. Efficient four fragment cloning for the construction of vectors for targeted gene replacement in filamentous fungi. BMC Mol. Biol. 9, 70 (2008).

137. Santhanam, P. Random insertional mutagenesis in fungal genomes to identify virulence factors. in 509–517 (2012). doi:10.1007/978-1-61779-501-5_31.

138. Bradford, M. M. A rapid and sensitive method for the quantitation of microgram quantities of protein utilizing the principle of protein-dye binding. Anal. Biochem. 72, 248–254 (1976).

139. Gabriel, L., et al. BRAKER3: Fully automated genome annotation using RNA-seq and protein evidence with GeneMark-ETP, AUGUSTUS, and TSEBRA. Genome Res. 34, 769–777 (2024).

140. Depotter, J. R. L., et al. The interspecific fungal hybrid *Verticillium longisporum* displays subgenome-specific gene expression. mBio 12, (2021).

141. Keilwagen, J., Hartung, F., Paulini, M., Twardziok, S. O. & Grau, J. Combining RNA-seq data and homology-based gene prediction for plants, animals and fungi. BMC Bioinformatics 19, 189 (2018).

142. Keilwagen, J., et al. Using intron position conservation for homology-based gene prediction. Nucleic Acids Res. 44, e89–e89 (2016).

143. Steinegger, M. & Söding, J. MMseqs2 enables sensitive protein sequence searching for the analysis of massive data sets. Nat. Biotechnol. 35, 1026–1028 (2017).

144. Xiong, W., He, L., Lai, J., Dooner, H. K. & Du, C. HelitronScanner uncovers a large overlooked cache of *Helitron* transposons in many plant genomes. Proceedings of the National Academy of Sciences 111, 10263–10268 (2014).

145. Hui, W., Gel, Y. R. & Gastwirth, J. L. lawstat: An R package for law, Public policy and biostatistics. J. Stat. Softw. 28, (2008).

146. Kabsch, W. XDS. Acta Crystallogr. D Biol. Crystallogr. 66, 125–132 (2010).

147. Vonrhein, C., et al. Data processing and analysis with the *autoPROC* toolbox. Acta Crystallogr. D Biol. Crystallogr. 67, 293–302 (2011).

148. Evans, P. R. & Murshudov, G. N. How good are my data and what is the resolution? Acta Crystallogr. D Biol. Crystallogr. 69, 1204–1214 (2013).

149. Winn, M. D., et al. Overview of the *CCP* 4 suite and current developments. Acta Crystallogr. D Biol. Crystallogr. 67, 235–242 (2011).

150. Evans, P. Scaling and assessment of data quality. Acta Crystallogr. D Biol. Crystallogr. 62, 72–82 (2006).

151. Skubák, P. & Pannu, N. S. Automatic protein structure solution from weak X-ray data. Nat. Commun. 4, 2777 (2013).

152. Emsley, P. & Cowtan, K. *Coot*: model-building tools for molecular graphics. Acta Crystallogr. D Biol. Crystallogr. 60, 2126–2132 (2004).

153. Murshudov, G. N., et al. *REFMAC* 5 for the refinement of macromolecular crystal structures. Acta Crystallogr. D Biol. Crystallogr. 67, 355–367 (2011).

