## Supplemental Figures 1-17 for "A structurally unique effector shared between *Verticillium dahliae* and *Fusarium oxysporum* is involved in cotton defoliation and virulence on other hosts"

<u>**CONTENT:**</u>

**Supplemental Figures 1 – 17**

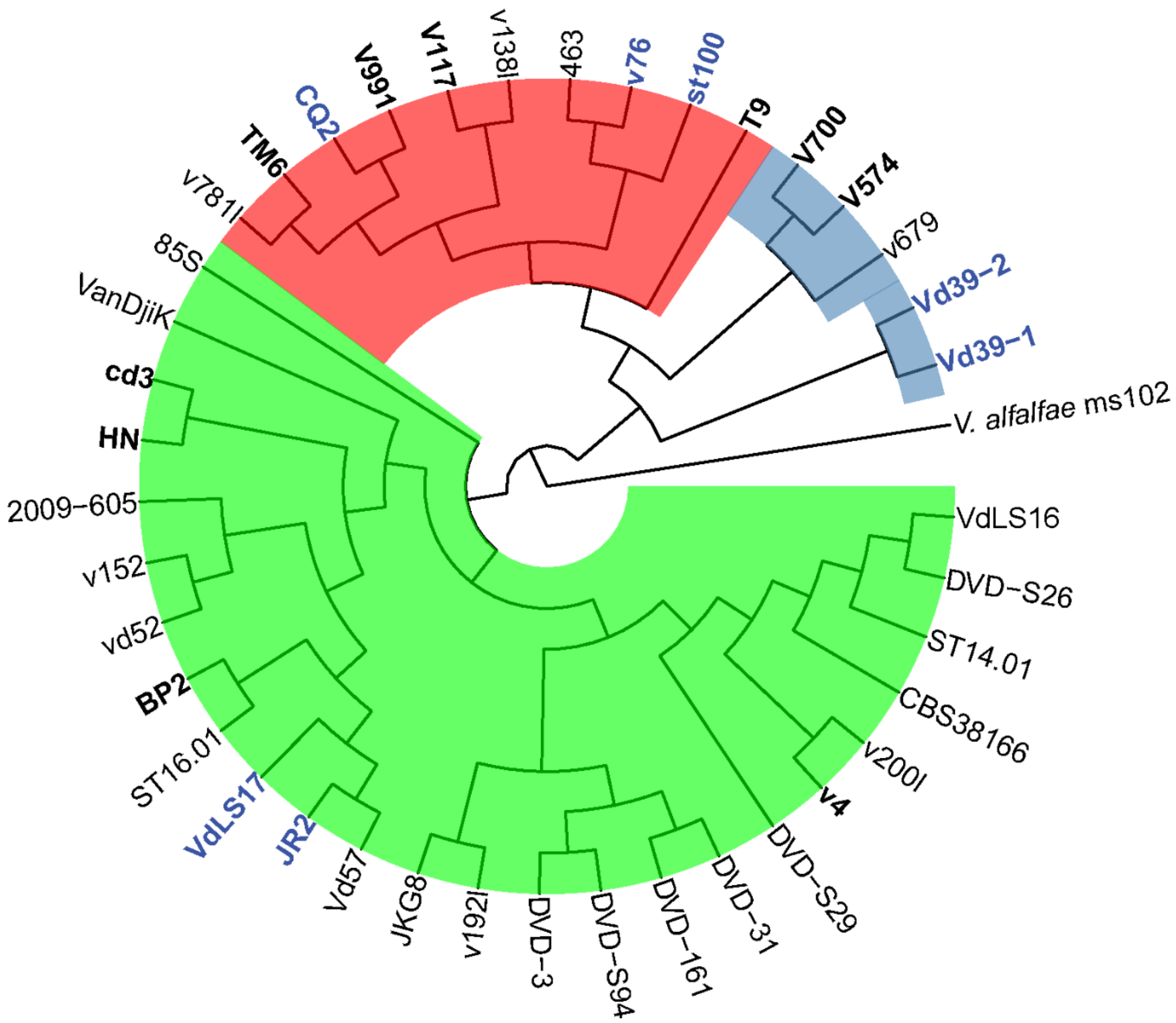


**Figure S1. Phylogenetic tree of in house-sequenced *V. dahliae* strains.** *V. dahliae* strains clustering with known D pathotype strains (TM6, V991, V117 and T9) are showed in lightred, while strains clustering with known ND pathotype strains (V4, BP2, cd3 and HN) are showed in light-green. Strains that distinct from the D and ND pathotype groups are showed in light-blue. Strains that used for phenotypic characterization in this work are display in bold with blue color. Phylogenetic relationship between sequenced *V. dahliae* strains was inferred using RealPhy (Bertels et al., 2014) and *V. dahliae* strain JR2 used as reference. *V. alfalfae* ms102 was used as root of the tree.

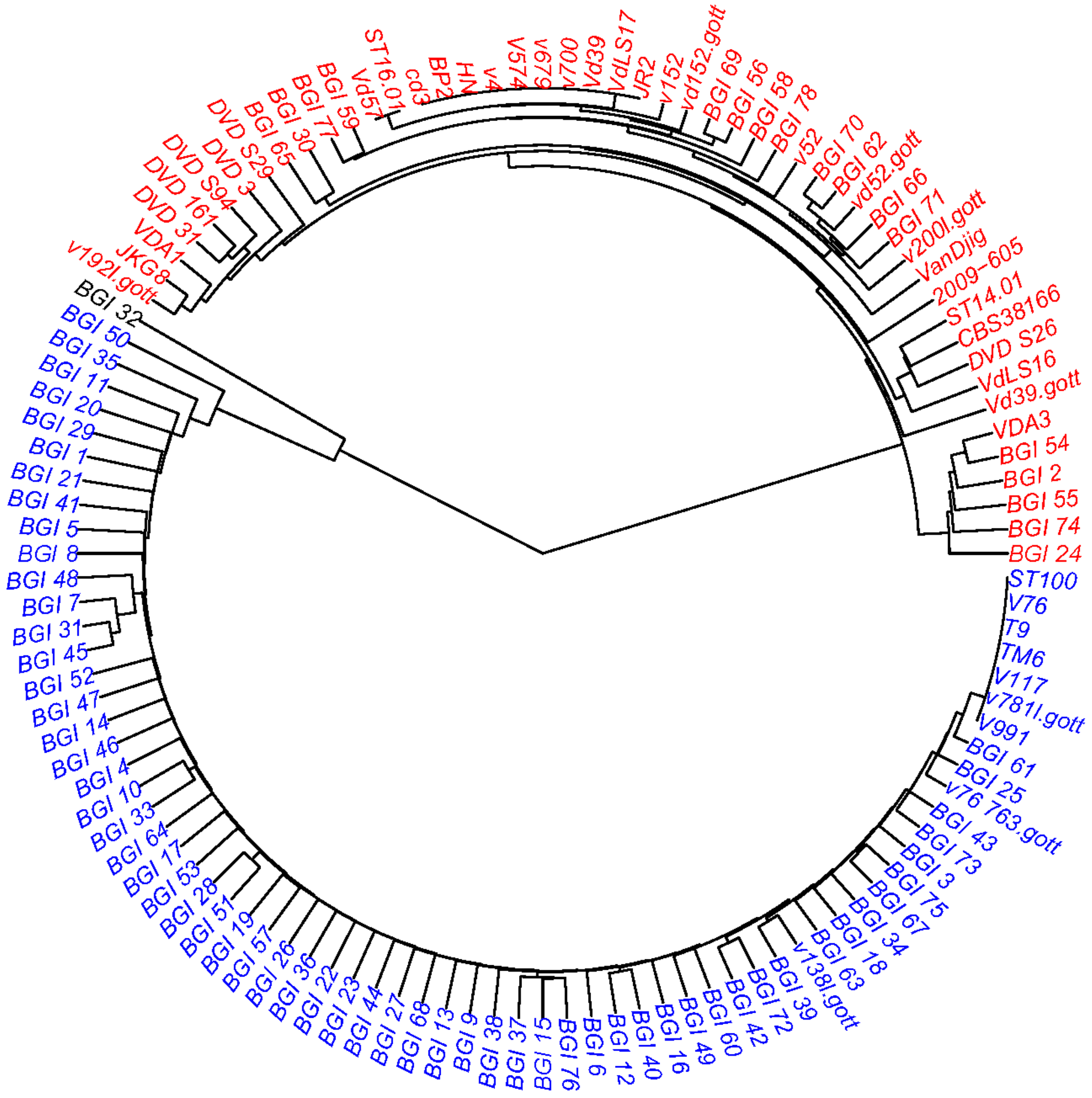


**Figure S2. Clustering analysis of seventy-two sequenced *V. dahliae* strains.** Sequences of seventy-two *V. dahliae* strains were aligned on the assembled genome of CQ2 and clustered in three groups based on presence/absence polymorphism. Strains clustering with known D pathotype strains are displayed in blue, while strains clustering with ND pathotype strains are showed in red. Strain BGI_32 that is the most divergent from the two groups showed in black.

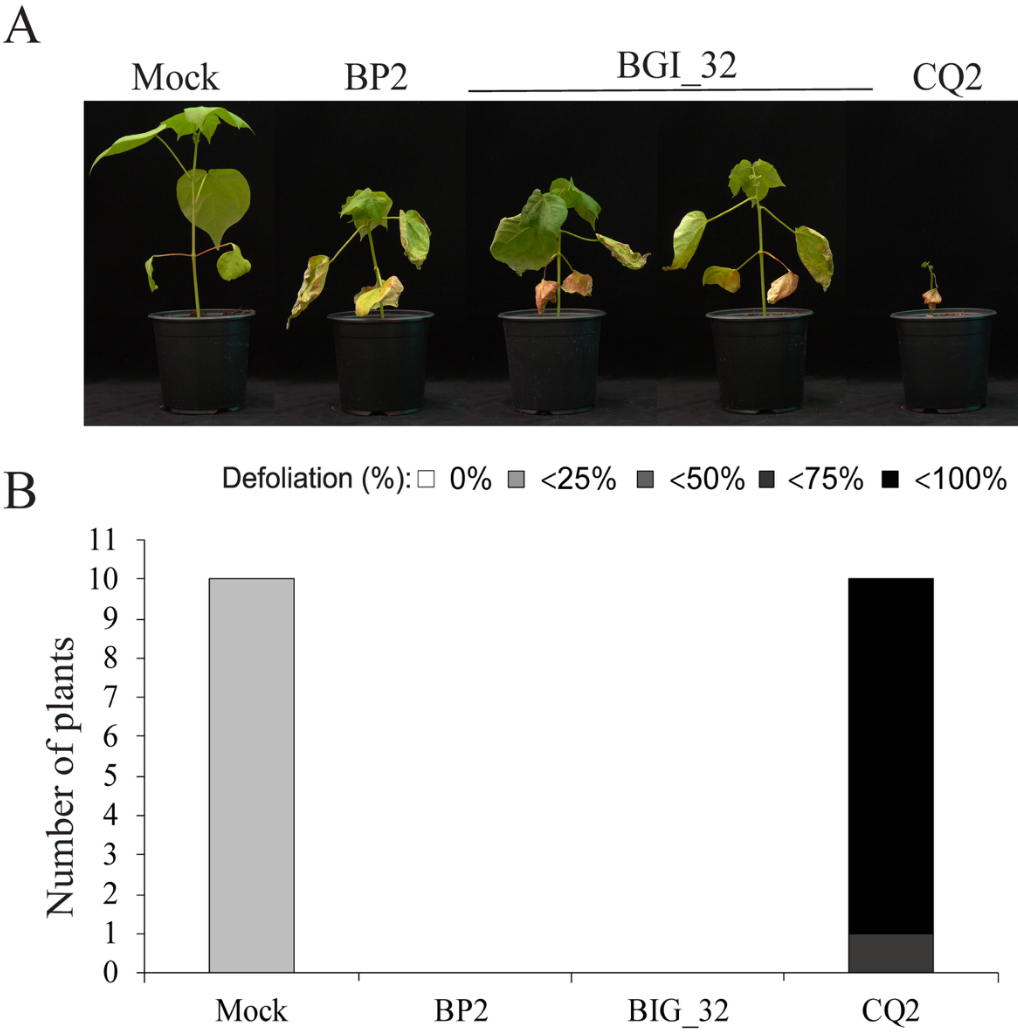


**Figure S3. Phenotype of cotton plants inoculated with *V. dahliae* strain BGI32.** (A) Typical phenotype of cotton plants (cv. Xinluzao63) upon mock-inoculation or inoculation with BP2, BGI32 and CQ2 at 28 days post inoculation (dpi). ND pathotype strain BP2 and D pathotype strain CQ2 were used as inoculation controls. (B) Defoliation was classified as 0 (0% leaf drop off), 1 (<25% leaf drop off), 2 (<50% leaf drop off), 3 (<75% leaf drop off) and 4 (<100% leaf drop off). Inoculation experiments were performed with ten plants for each fungal strain and repeated twice independently with similar results.

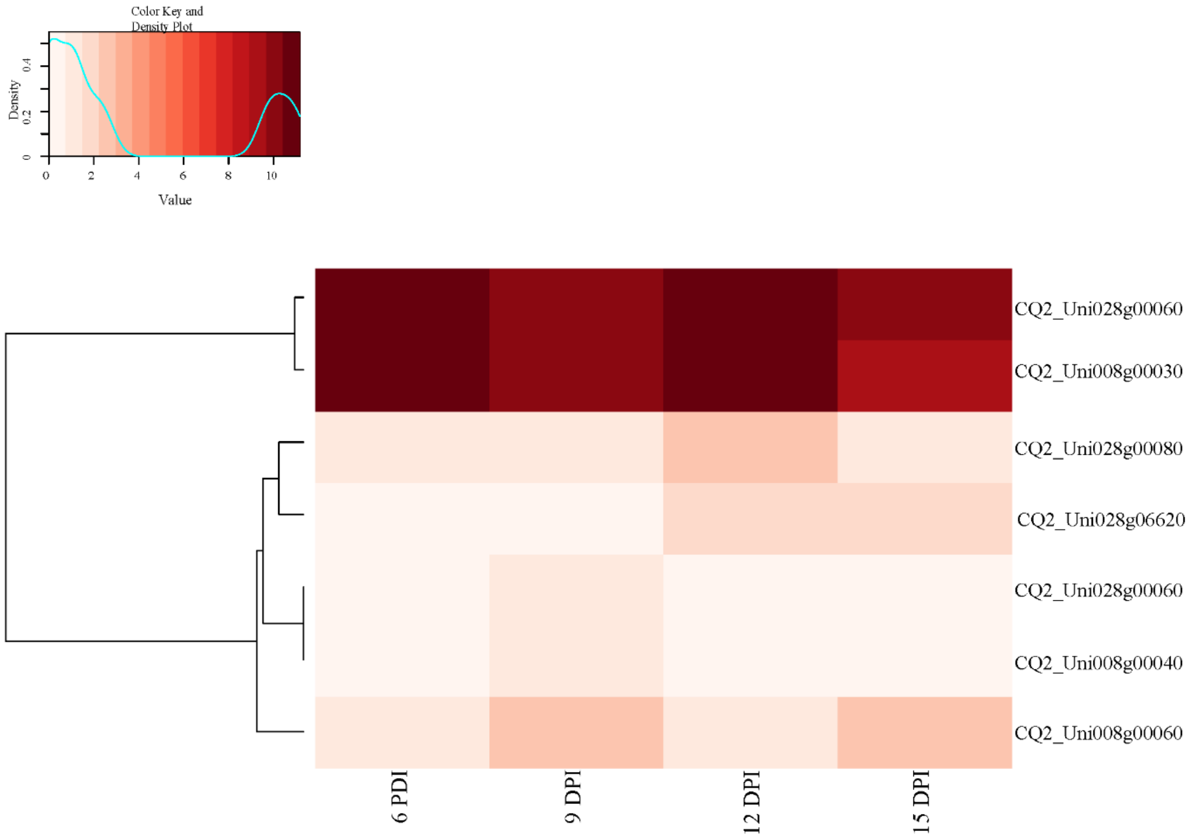


**Figure S4. Expression analysis of the seven candidate genes.** The heatmap showed the expression level of each gene during a time course of cotton infected by D pathotype strain V991 at 6, 9, 12 and 15 days post inoculation (DPI). The scaled expression values are color-coded according to the scale bar in the left top corner.

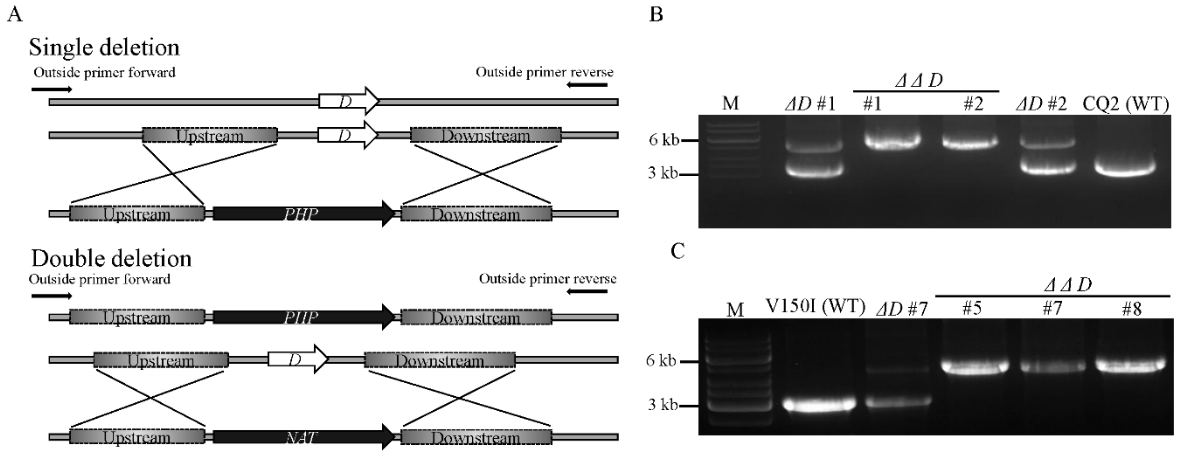


**Figure S5. Construction and verification of *D* single and double deletion mutants.** (A) Schematic representation of the homologous recombination events to establish targeted replacement of *D* gene with phosphotransferase (HPH) and the nourseothricin resistance gene cassette (NAT). (B) Verification of D single deletion (Δ*D*) and double deletion mutants (ΔΔ*D*) in *V. dahliae* cotton defoliating strain CQ2 by PCR. Amplicons generated with outside primers indicated in panel A are shown for wild type strain CQ2 (WT), two Δ*D* mutants (#1 and #2) and two ΔΔ*D* mutants (#1 and #2). (C) Verification of *D* single deletion (Δ*D*) and double deletion mutants (ΔΔ*D*) in *V. dahliae* olive defoliating strain V150I by PCR. Amplicons generated with outside primers indicated in panel A are shown for wild-type strain V150I (WT), one Δ*D* mutant (#7) and three ΔΔ*D* mutants (#5, #7 and #8).

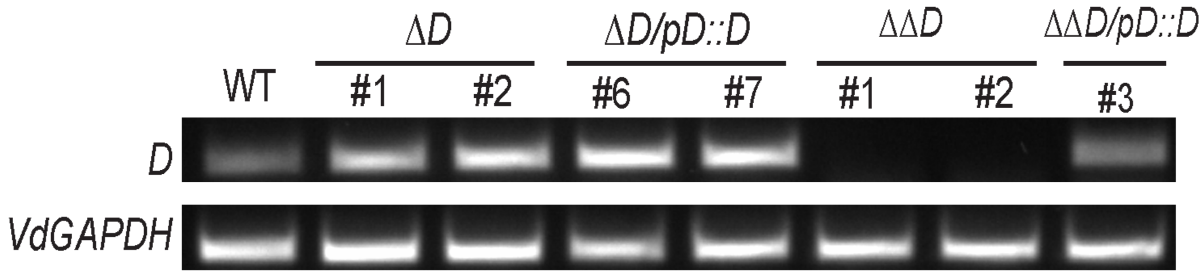


**Figure S6. Detection of *D* gene transcripts in various *D* deletion and complementation strains.** Amplification of *D* gene fragment (from left to right) from cDNA in wild type strain CQ2 (WT), two Δ*D* strains (#1 and #2), two Δ*D* complementation strains (Δ*D*/p*D::D* #1 and #2), two ΔΔ*D* strains (#1 and #2) and one ΔΔ*D* complementation strain (ΔΔ*D*/p*D*::*D* #3). *V. dahliae GAPDH* gene was used as endogenous control.

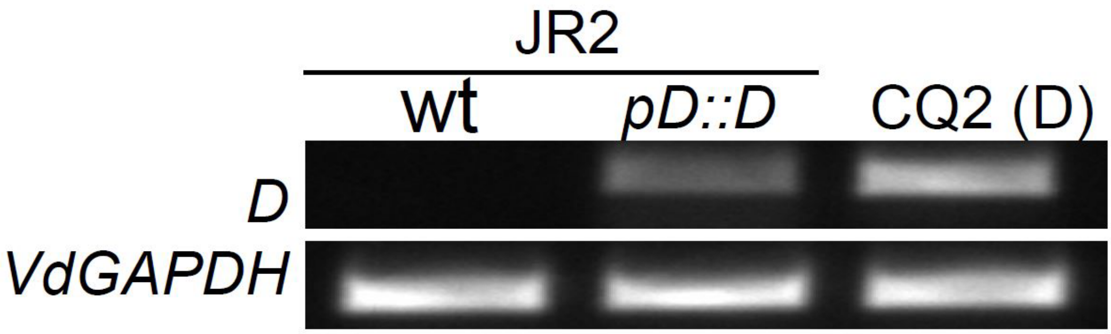


**Figure S7. Expression of *D* gene (p*D*::*D*) in ND pathotype strain JR2.** Amplification of *D* gene fragment (from left to right) from cDNA in wild type strain JR2, one *D* expression (p*D::D*) transformant of JR2 and CQ2 (used as positive control). *V. dahliae GAPDH* gene was used as endogenous control.

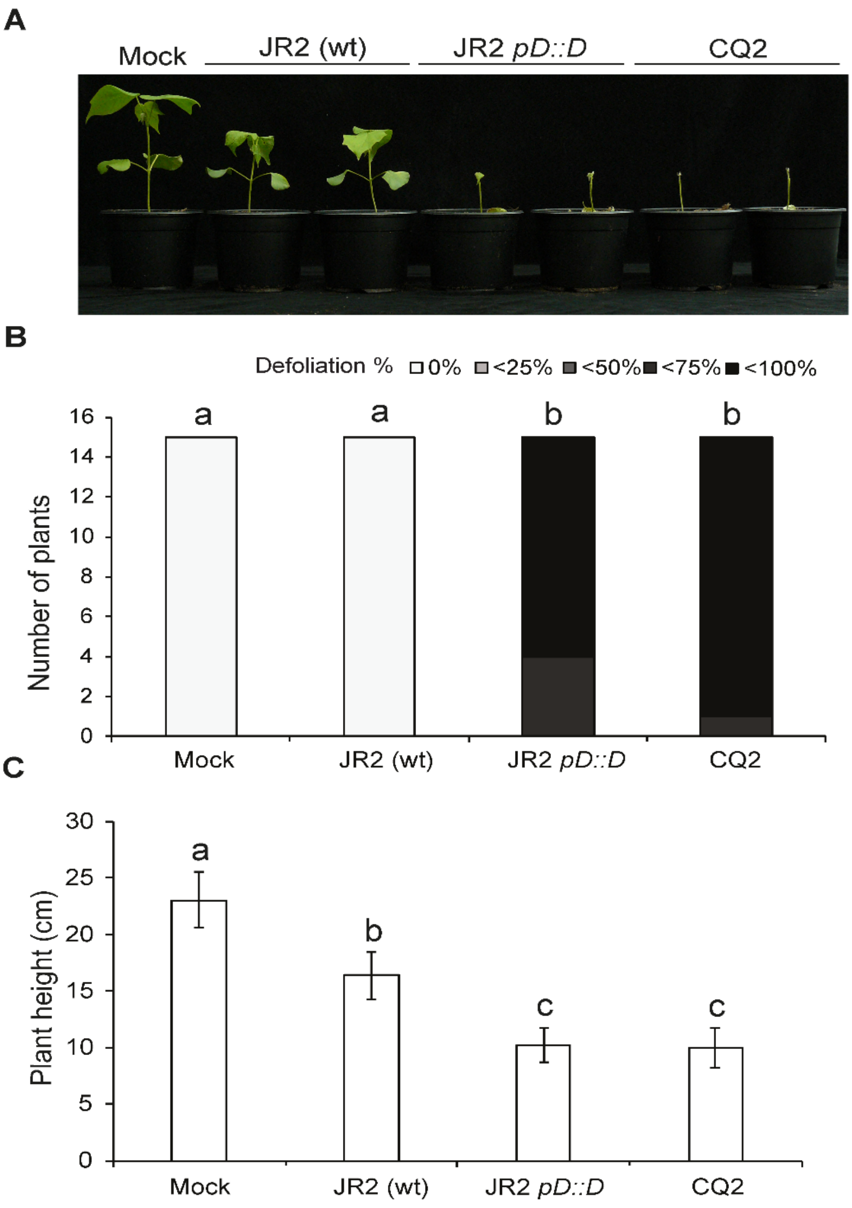


**Figure S8. Introduction of *D* gene in ND pathotype strain results in defoliation symptoms.** (A) Typical phenotype of cotton (cv. Xinluzao63) plants that were mock-inoculated or inoculated with ND pathotype strain JR2, one *D* expression transformant of JR2 (JR2 p*D*::*D*) and D pathotype strain CQ2 at 28 days post inoculation (dpi). (B) Defoliation was classified as 0 (0% leaves drop off), 1 (<25% leaves drop off), 2 (<50% leaves drop off), 3 (<75% leaves drop off) and 4 (<100% leaves drop off) at 28 dpi. (C) Fungal biomass as determined with real-time PCR at 28 dpi. Bars indicate the *V. dahliae* biomass relatively to the cotton biomass. Significant differences were calculated with the Mann-Whitney U test ($P < 0.05$) and depicted by different letter labels. Error bars represent the standard error.

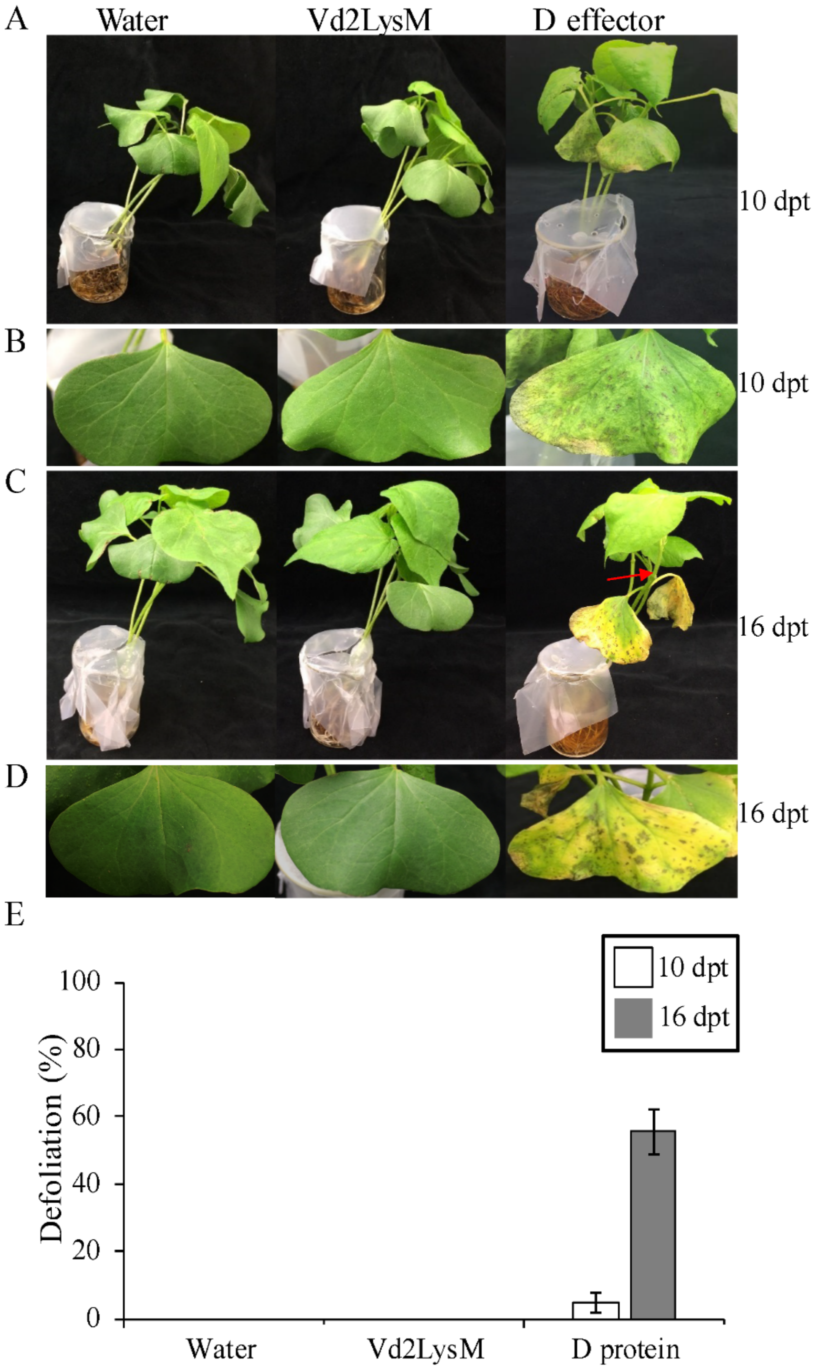


**Figure S9. D effector protein induces cotton defoliation *in vitro*.** (A-D) Typical appearance of cotton seedlings (cv. Xinluzao63) treated with D effector protein at 10 days (A-B) and 16 days (C-D). Note, marginal chlorosis and wilting symptoms on cotyledons at 10 days (B) and severe chlorosis, and leaf drop off (red arrow; D). (E) Bars represent the average defoliation of two biological replicates with standard deviation. Experiments were repeated twice independently with similar results.

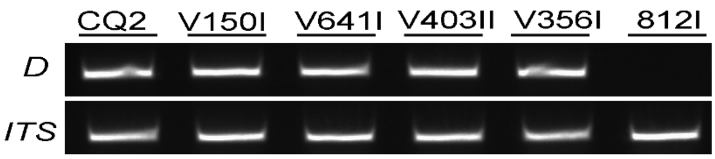


**Figure S10. PCR detection of *D* gene in *V. dahliae* strains.** Amplification of *D* gene fragment (from left to right) from genomic DNA in CQ2, V150I, V641I, V403I, V356I, 812I. As an endogenous control, a fragment of the *Verticillium* ITS region was amplified.

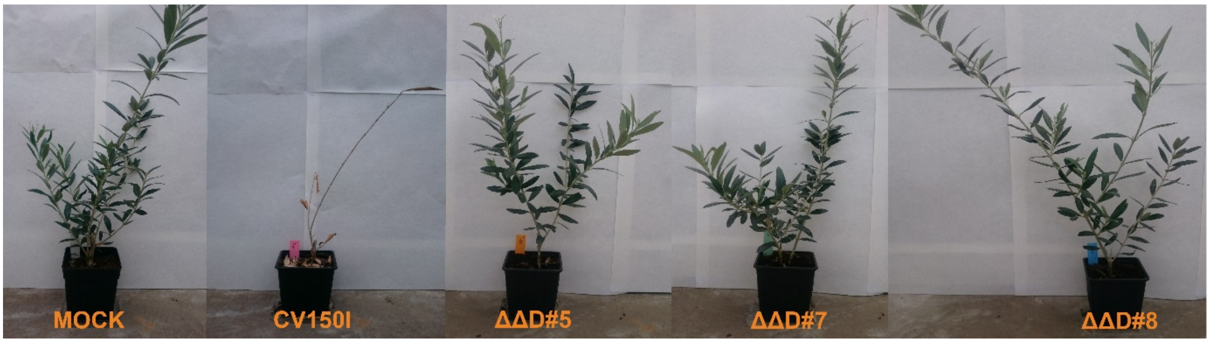


**Figure S11. The D effector is responsible for olive defoliation.** Typical phenotype of olive (cv. Picual) upon mock-inoculation or inoculation with wild type strain CV150I (WT), and three ΔΔ*D* mutants (ΔΔ*D* #5, ΔΔ*D* #7 and ΔΔ*D* #8) at 132 days post inoculation (dpi).

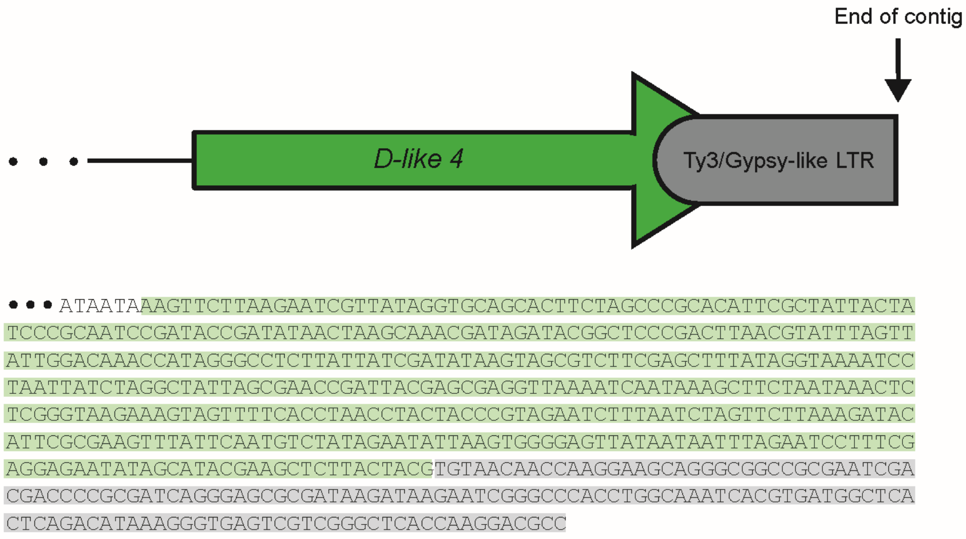


**Figure S12. The *D-like 4* allele is disrupted by a retrotransposon of the family Ty3/Gypsy-like**. Schematic representation of the *D-like4* allele, showing the interruption of its sequence (represented in green) by a retrotransposon of the family Ty3/Gypsy-like (represented in grey). Only a short portion (174 bp) of the retrotransposon is present before the end of the contig in which the D-like 4 allele is present in both *V. dahliae* strains that carry this allele (V574 and V700).

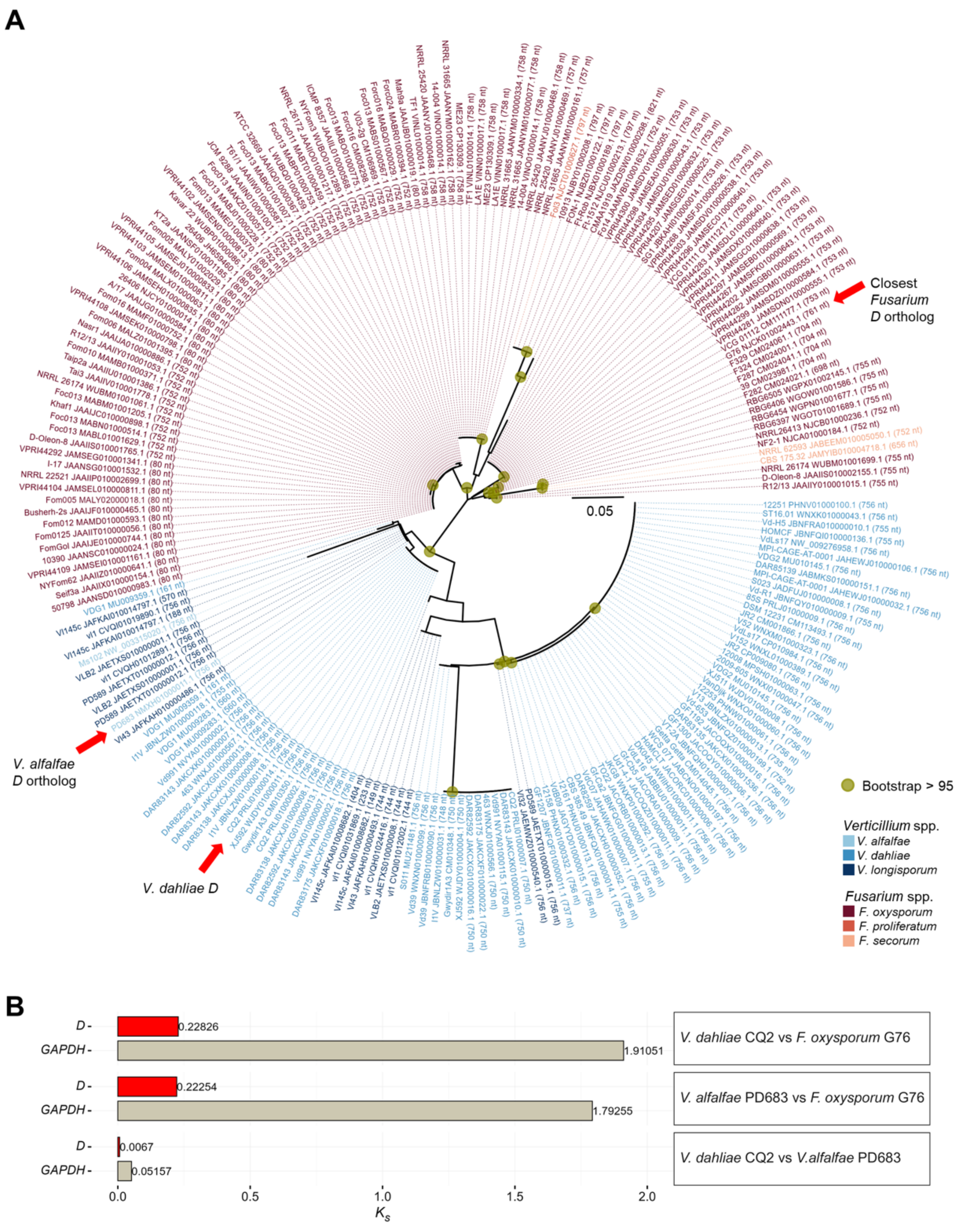


**Fig. S13. *D* gene horizontal transfer.** (A) Maximum-likelihood phylogenetic tree of *D* homologs inferred using a nucleotide substitution model. Scale bars indicate nucleotide substitutions per site. Circles at nodes denote bootstrap support values >95% based on 1,000 replicates. (B) Rates of synonymous substitutions per synonymous site ($K_s$) between species. The *D* homologs selected for testing are indicated by red arrows in (A). Glyceraldehyde 3-phosphate dehydrogenase (*GAPDH*) genes from the same strains were used as vertically inherited core housekeeping gene references.

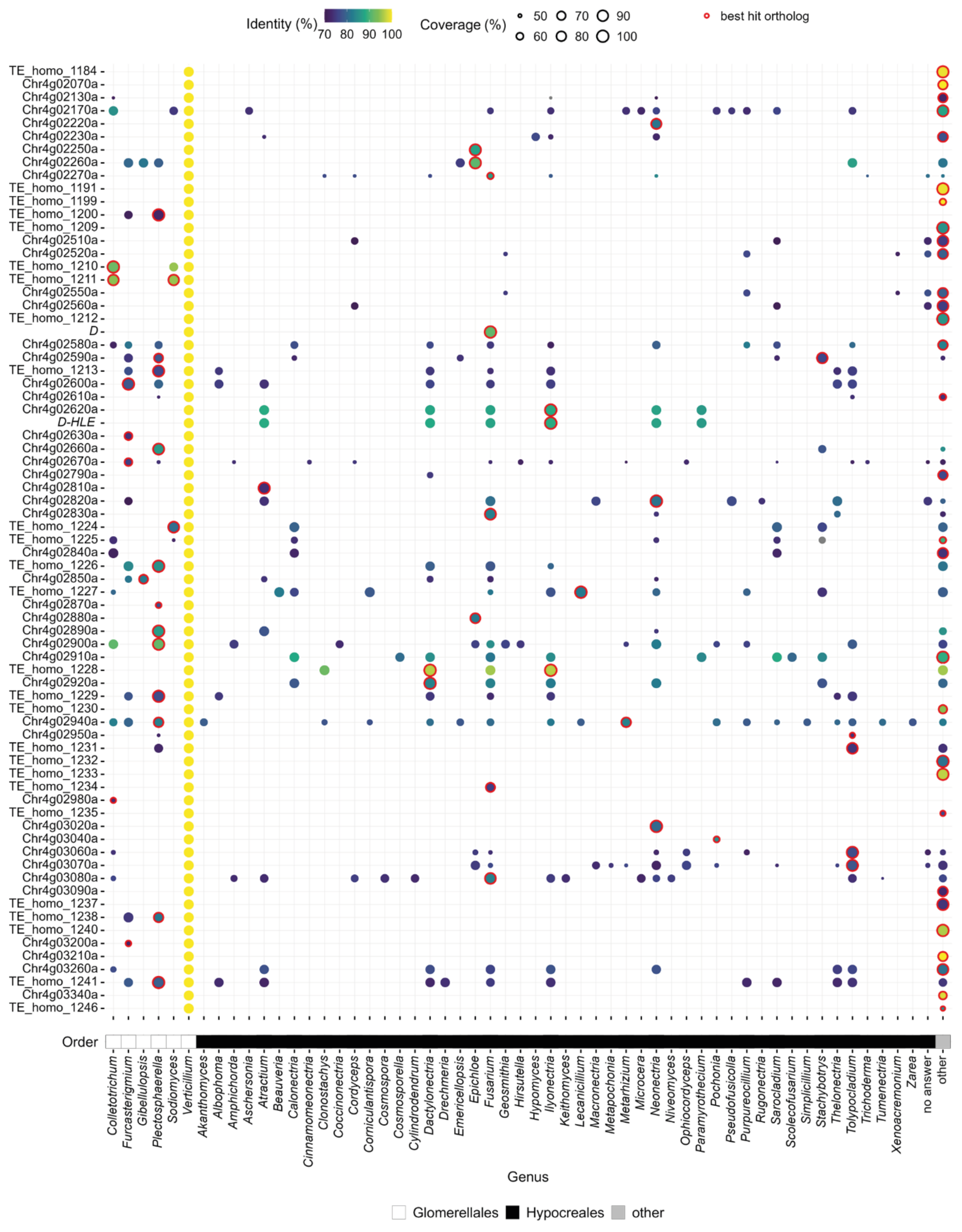


**Fig. S14. Distribution of homologs of *Ar1h2 Starship* cargo genes and transposable elements (TEs) across Pezizomycotina.** Cargo elements from the *Ar1h2 Starship* in *V. dahliae* strain CQ2 were used as queries. Circles indicate the coverage and sequence identity of the best hits in each genus within the orders Glomerellales and Hypocreales, as well as in all other genera. Best hits across all genera excluding *Verticillium* are outlined in red.

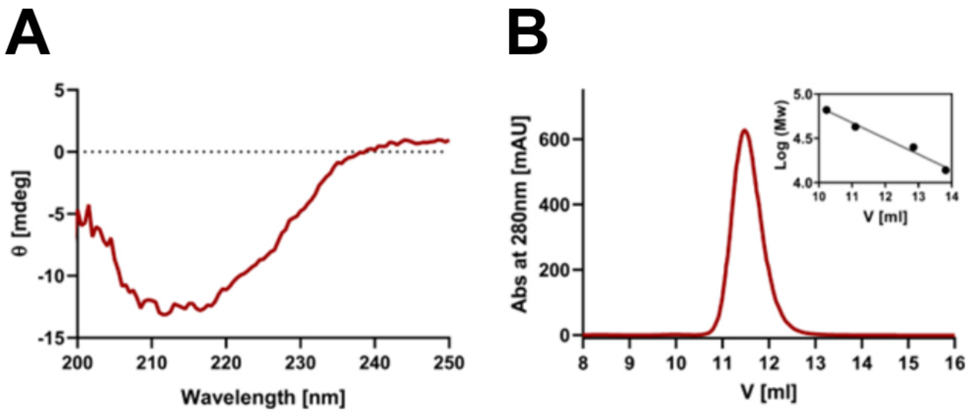


**Figure S15. Circular dichroism spectroscopy and size exclusion chromatography analysis of the D*Fov* protein.** (A) Far-UV CD spectra of *E. coli*-produced D*Fov* protein (0.2 mg/ml) in 20 mM Tris-HCl pH 7.5, 150 mM NaCl buffer. The spectrum reveals that the recombinant protein is well folded with a prevalently beta-sheet secondary structure. (B) Size exclusion chromatographic elution profile of purified D*Fov*. The protein, loaded on a Superdex 75 10/300 column, eluted as a single sharp peak at 11.7 ml, corresponding to an apparent molecular weight of 35 kDa. Considering that the purified protein, including the affinity tag, has a predicted Mw of 26.6 kDa, this value indicates that D*Fov* is monomeric and displays an elongated shape. *Inset*: the column was calibrated by running the following protein standards: albumin (66.5 kDa), ovalbumin (43 kDa), chymotrypsinogen (25 kDa) and ribonuclease A (13.7 kDa).

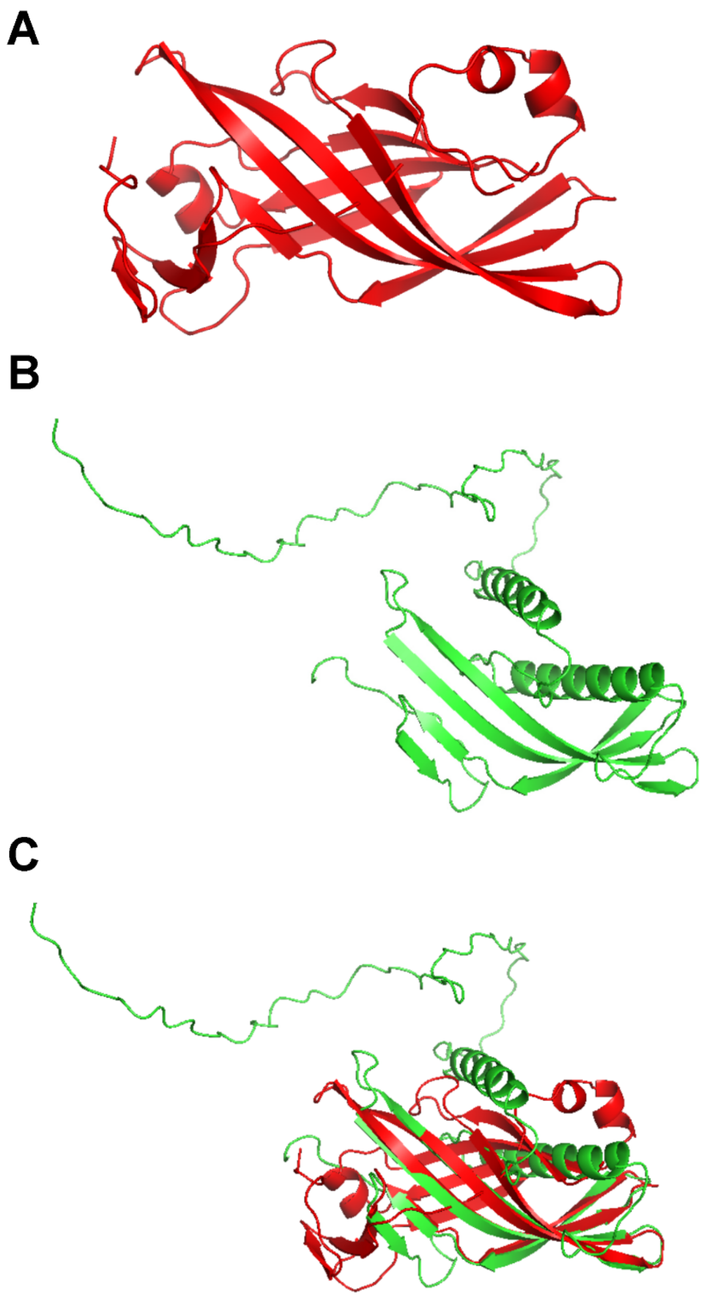


**Figure S16. The AlphaFold3-predicted D*Fov* model diverges from the D*Fov* protein structure determined with X-ray crystallography.** (A) D*Fov* crystal structure, determined by X-ray crystallography. (B) D*Fov* model predicted with AlphaFold3. (C) Structural alignment of the D*Fov* crystal structure and the AlphaFold3 model. TM-score indicates topological similarity between two protein structures reported on a scale from 0 to 1, where 1 indicates a perfect match, while the RMSD score indicates the average deviation between the corresponding atoms of two proteins with values lower than 2 indicating significant similarity between structures.

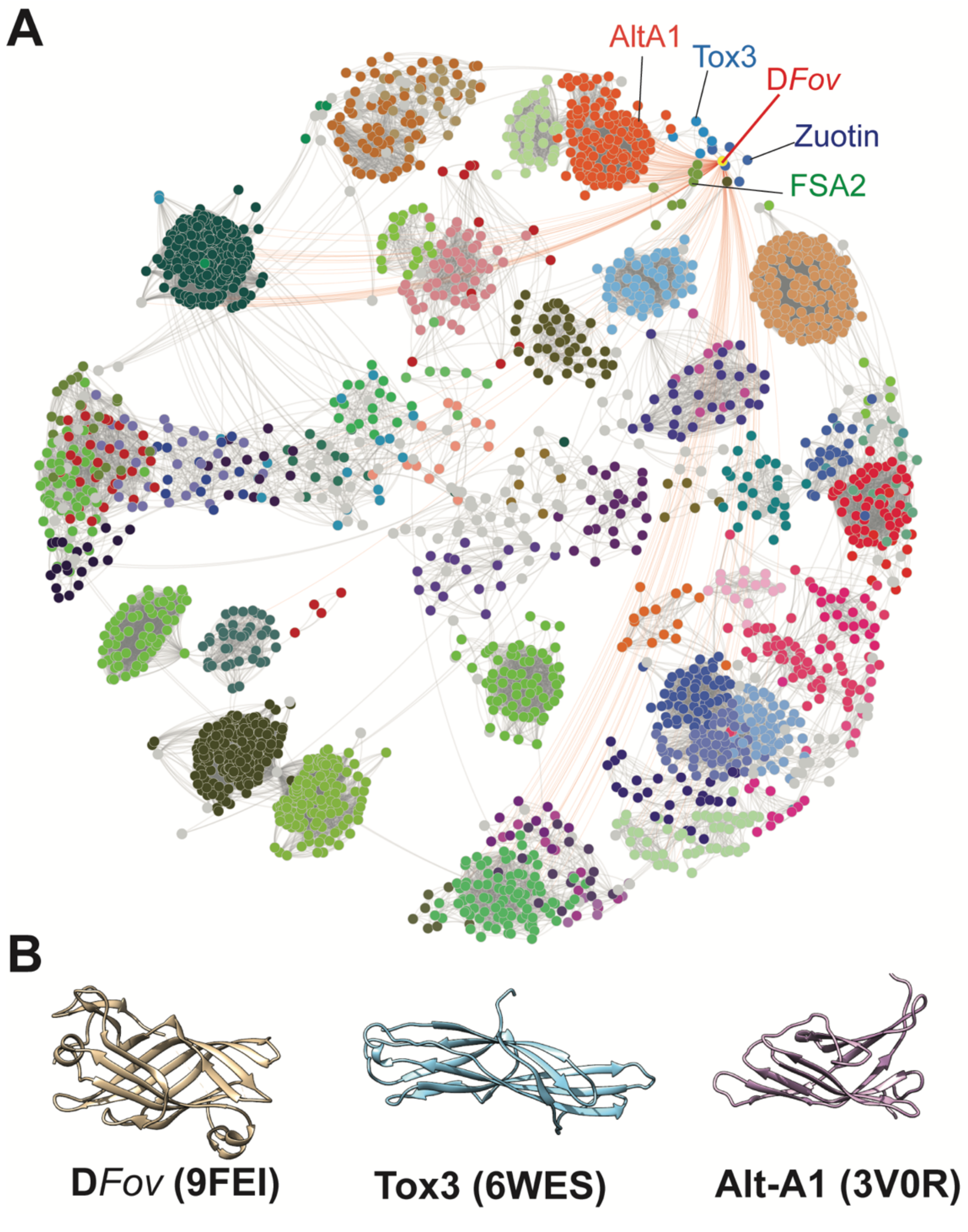


**Figure S17. D*Fov* is a structurally unique effector.** (A) Structural similarity network based on pairwise DALI Z-scores. Each dot represents a protein structure while different colors represent different structural families. The D*Fov* protein structure is indicated with as well as effector proteins that are somewhat similar, including Alt-A1, Tox3, Zuotin and FSA2. (B) D*Fov*, Tox3 and Alt-A1 protein structures.
